# STAG2 cohesin cooperates with DREAM to maintain quiescence and suppress tumourigenesis in the urothelium

**DOI:** 10.1101/2025.10.10.681713

**Authors:** Maria Ramal, Eleonora Lapi, Mark Kalisz, Jaime Martínez de Villarreal, Meritxell Novillo-Font, Miriam Marqués, Eduardo Zarzuela, Marta Isasa, Osvaldo Graña-Castro, Jianming Xu, Ana Cayuela, François Radvanyi, Mireia Novell, Jorge L. Martínez-Torrecuadrada, María J. Barallobre, Isabelle Bernard-Pierrot, Maria Marti-Marimon, Maria L. Arbonés, Marc A. Marti-Renom, Francisco X. Real

## Abstract

The maintenance of quiescence is essential for tissue homeostasis. *STAG2* is one of the few genes mutated in the normal urothelium of organ donors, with mutant cells undergoing positive selection ^1^. *STAG2* is also a major tumour suppressor gene ^2–4^ and its inactivation is an early event in bladder carcinogenesis ^1,3^. However, how STAG2, a cohesin component, regulates urothelial homeostasis remains largely unknown. Here, we demonstrate that *Stag2* inactivation in normal murine urothelial cells interferes with differentiation programs, triggers transient cell cycle entry, and primes cells for clonal expansion under stress. Moreover, STAG2 loss enhances tumor formation in urothelial cells expressing mutant *FGFR3* - the key oncogene in bladder cancer ^5^. We reveal that STAG2 cooperates with the DREAM transcriptional complex, a master regulator of quiescence ^6,7^, by binding to shared genomic sites, including cell cycle control genes. STAG2 loss alters DREAM target expression, complex composition, and chromatin distribution, and leads to rewiring of chromatin interactions involving DREAM binding motifs in genes critical for cell cycle entry. Our findings provide compelling evidence that STAG2 loss disrupts in 3D genome organization through a novel mechanism involving the DREAM complex, thereby impairing homeostatic quiescence and increasing oncogenic sensitivity.

## Introduction

The maintenance of quiescence is critical for homeostasis and to avoid tumourigenesis in many tissues. Evading growth suppression and sustaining proliferative signaling are two major hallmarks of cancer and classical oncogenes and tumour suppressor genes play key roles in these processes ^8^. How quiescence is regulated and maintained, or perturbed, remains an important question to understand tumour initiation.

Recently, the occurrence of somatic mutations in normal tissues has revealed critical and complex roles of mutational processes in homeostasis and disease ^1,9–11^. Mutations can confer a selective growth advantage to cell clones, but we still have a poor understanding of the mechanisms involved and their consequences. Clonal hematopoiesis, which is associated with an increased risk of hematopoietic neoplasms and other conditions, has been most extensively studied ^12,13^. However, less is known about clonal dynamics in other tissues ^14–16^. In the normal urothelium of organ donors, mutations were identified in some bladder cancer drivers, but not in others, indicating selective, stage-specific, roles of cancer genes in disease ^1,9^. The significance of such mutations remains to be determined.

*STAG2*, which encodes a cohesin subunit, is one of the genes commonly mutated in the urothelium of healthy subjects and its inactivation is associated with positive selection ^1^. Cohesin plays important roles beyond chromosome segregation, including 3D genome organization, gene regulation, and DNA repair ^17,18^. Bladder cancer is the tumour with the highest frequency of *STAG2* alterations. The majority of *STAG2* mutations lead to reduced mRNA and protein expression and result in loss-of-function ^3,19^. Low-grade bladder cancers are enriched in *STAG2* alterations, which are significantly associated with oncogenic *FGFR3* mutations ^2,3^. These tumours are generally indolent but can become aggressive upon the acquisition of additional genetic alterations (e.g., *CDKN2A*, *TP53*). *STAG2* mutations occur in tumours lacking aneuploidy, suggesting roles beyond chromosome segregation ^2,3^.

Here, we leverage genetic mouse models, organoids, and genomic tools to investigate the impact of *Stag2* inactivation on urothelial homeostasis and demonstrate that it plays a critical role in maintaining epithelial quiescence. We show that STAG2 cooperates with DREAM to control the expression of late cell cycle genes. The DREAM complex is a transcriptional regulator consisting of dimerization partner (DP), RB-like 1 (RBL1/p130), E2F4, and the MuvB core, which represses cell cycle genes ^7,20^. Cyclin-CDK-mediated phosphorylation of p130/107 triggers DREAM disassembly, causing MuvB to dissociate from the repressor components p130 and E2F4 and to bind MYB-B and FOXM1 (DREAM/MuvB-MYB-B-FOXM1), thereby activating gene expression and promoting cell proliferation ^21–23^. Despite the crucial role of DREAM in cell cycle control, its physiological role and its involvement in cancer remain largely unexplored. We show that STAG2 loss results in changes in DREAM composition, expression of its target genes, and in chromatin looping involving DREAM binding motifs. These findings reveal a novel mechanism by which STAG2 and DREAM cooperate to up-regulate cell cycle gene expression, providing a framework for understanding their role in urothelial homeostasis and how *STAG2* inactivation sensitizes cells to oncogenic events.

## Results

### *Stag2* deficiency primes urothelial cells for proliferation

To investigate the effects of *Stag2* inactivation on urothelial homeostasis *in vivo*, we generated *Tg.UBC^CreERT^*^2^*;Stag2^lox/lox^;Rosa26^mT/mG^*mice (hereafter referred to as *Stag2*-KO mice once recombination has been induced) ^24^. Adult mice were fed a tamoxifen (TMX)-containing diet for one week and were sacrificed at one week and at one month (**Suppl. Fig. 1a**). We confirmed that STAG2 was lost in >95% of cells in the bladder of *Stag2-*KO mice at both time points (**Fig. 1a, Suppl. Fig. 1b,c**). There were no major histological changes and expression of urothelial (UPK3A) and basal (KRT5 and TP63) lineage markers was similar in control and *Stag2*-KO urothelial cells at both time points (**Fig. 1a, Suppl. Fig. 1c**). In contrast, we observed a significant increase in KI67 expression (*P*=0.022) and BrdU incorporation (*P*=0.049) in *Stag2*-KO mice at one week that was no longer detectable at one month (**Fig. 1a,b, Suppl. Fig. 1c**). We also found a significant increase of gH2AX in *Stag2*-KO urothelial cells (**Suppl. Fig. 1d**), suggestive of enhanced DNA damage response, but the proportion of cleaved caspase 3^+^ and p21^+^ cells was similar in control and recombined urothelia (**Suppl. Fig. 1e**). These results indicate that the sole inactivation of *Stag2* in basal conditions triggers a transient cell cycle entry response but does not alter urothelial architecture.

**Figure 1.**
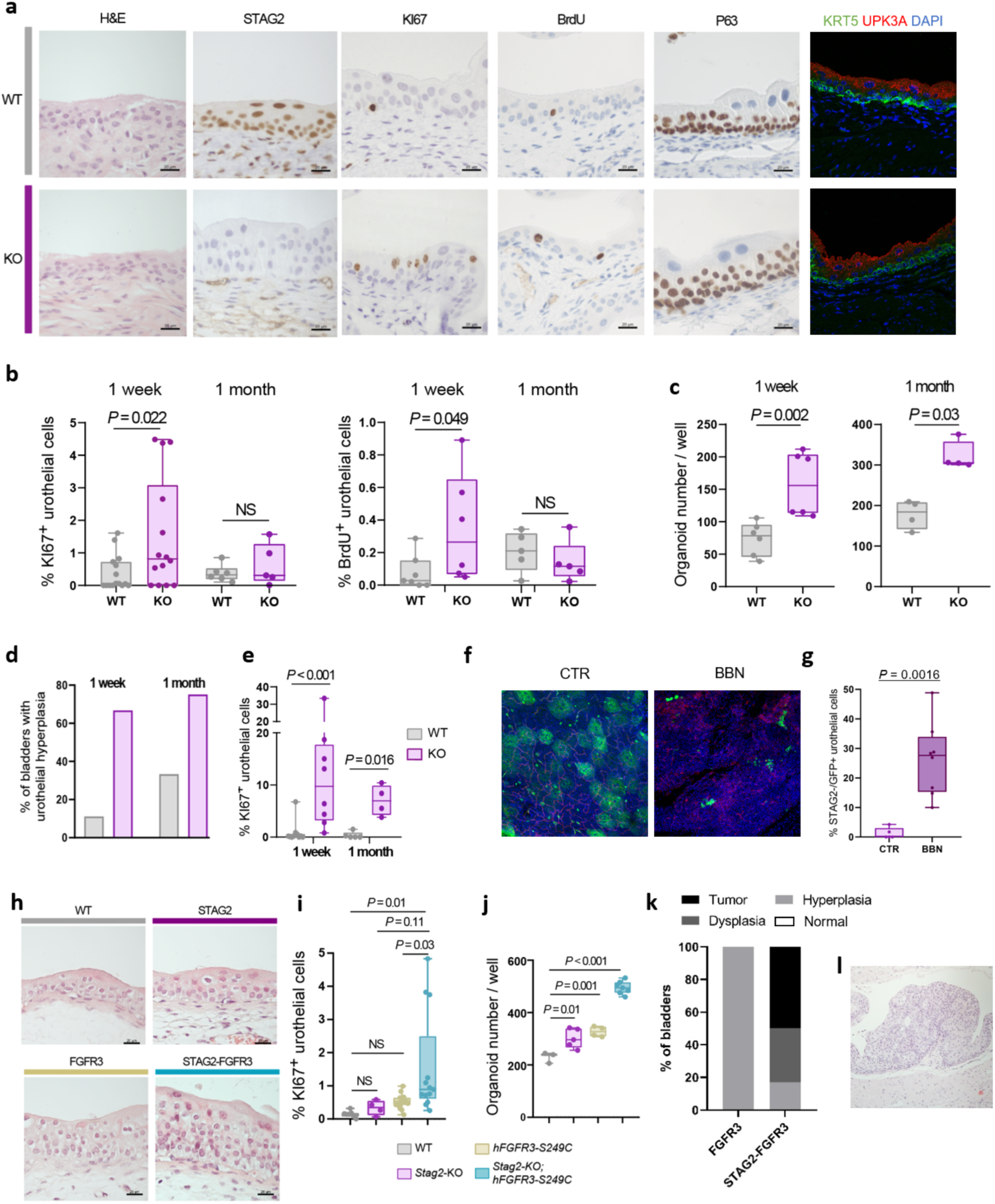
*Stag2* inactivation primes urothelial cells for proliferation. **a-g,** 8-12 week-old *Tg.hUBC-CreERT2;Stag2^lox/lox^;Rosa^mTmG^* mice received TMX for one week and the urothelium was analyzed at one week or one month. **a**, H&E, IHC for STAG2, KI67, BrdU, and p63, and immunofluorescence for KRT5 and UPK3A in WT and *Stag2-*KO bladder sections. Scale bar, 20 μm. **b**, Increased proliferation of *Stag2-*KO urothelial cells, assessed by KI67 expression and BrdU incorporation, at one week, but not at one month. Each data point represents an individual mouse. Statistical test: t-test for KI67 (normal distribution) and two-tailed Mann-Whitney for BrdU. **c**, Increased organoid-forming capacity of GFP^+^ urothelial cells from control and *Stag2*-KO mice at one week or one month after initiation of TMX administration. Each dot represents the mean value per organoid culture, established from an independent mouse bladder. Statistical test: two-tailed Mann-Whitney; one representative experiment. **d**, Percentage of WT and *Stag2-*KO bladders displaying hyperplasia at one week (n = 6-9 mice/group) and one month (n = 4-6 mice/group). **e**, Increased proliferation of *Stag2-*KO urothelial cells one week and one month after CYC administration. Each data point represents an individual mouse. Statistical test: two-tailed Mann-Whitney. **f**, Whole-mount urothelium of *Tg.hUBC-CreERT2;Stag2^lox/lox^* or *Stag2^lox/Y^*mice treated with low-dose TMX showing the number of GFP^+^ clones in basal conditions (CTR) and after 1 month of BBN administration. **g**, Quantification of STAG2^-^ GFP^+^ clones in basal conditions (CTR) and after 1 month of BBN treatment. Each data point represents an individual mouse. Statistical test: two-tailed Mann-Whitney. **h-l**, Mice were fed a TMX-containing diet over two weeks starting from weaning and sacrificed at 12 and 78 weeks of age. STAG2 loss cooperates with mutant FGFR3 to promote hyperplasia (**h**), urothelial cell proliferation (**i**), organoid-forming capacity (**j**), and dysplasia and tumour formation in aging mice (**i, j**; each data point represent an individual mouse. **k, l**; n = 6-8 mice/group). Representative H&E staining of a *Stag2*-KO*;hFGFR3-S249C* tumour. Statistical test: one-way ANOVA (Tukey’s multiple comparisons test; **i-j**).

The urothelium is one of the most quiescent epithelia but proliferation is readily induced upon tissue dissociation or injury ^25,26^. We quantified the organoid-forming capacity of freshly sorted urothelial cells from WT and KO mice in homeostatic conditions. *Stag2*-KO cells showed significantly higher organoid-forming capacity both 1 week and 1 month after TMX administration (**Fig. 1c**). Cyclophosphamide (CYC), a drug that selectively causes urothelial damage and regeneration ^27^, induced urothelial hyperplasia at 1 week and 1 month in *Stag2-* KO mice but not in control mice (**Fig. 1d**), consistent with the increased proliferation (**Fig. 1e**). These data support the notion that *Stag2*-KO urothelial cells have a higher potential for self-renewal. To extend these findings, we analyzed cell fate in *Stag2*-KO mice receiving N-butyl-N-(4-hydroxybutyl)-nitrosamine (BBN), a urothelium-specific carcinogen ^28^ (**Fig. 1f**). Recombination was induced with a low-dose of TMX, to ensure low-frequency labeling, and clonal density and size were analyzed via 3D-confocal imaging of the urothelium before and after one month of BBN exposure. A significant increase in STAG2^-^ GFP^+^ clones was observed in mice receiving BBN, indicating positive selection of mutant cells (*P* = 0.002; **Fig. 1g**).

In human bladder tumours, *STAG2* and *FGFR3* mutations show significant co-occurrence ^3^. We tested the cooperation of alterations in both genes using *UpkII_hFGFR3-S249C;Tg.Upk3a^CreERT^*^2^*;Stag2^lox/lox^; Rosa26^mT/mG^* mice fed over 2 weeks with a TMX-containing diet. At 12 weeks, *Tg.UpkII-hFGFR3-S249C* mice (n=15) showed mild thickening of the urothelium while 54% (n=13) *hFGFR3-S249C;STAG2*-KO mice showed hyperplasia (**Fig. 1h**), consistent with a significant increase of Ki67^+^ cells (**Fig. 1i**). At 78 weeks, 33% (n=6) of *hFGFR3-S249C;STAG2*-KO mice had urothelial dysplasia and 50% (n=6) had papillary tumours, while *hFGFR3-S249C* mice exhibited only hyperplasia (100%, n=8) (**Fig. 1k,l**). In addition, *hFGFR3-S249C;Stag2*-KO urothelial cells had significantly higher organoid-forming capacity compared to *Stag2*-KO or *hFGFR3-S249C* urothelial cells **(Fig. 1j**). Altogether, our findings suggest that STAG2 loss primes cells for proliferation and oncogenic transformation, highlighting the relevance of these tools to model human bladder carcinogenesis.

**Suppl. Figure 1.**
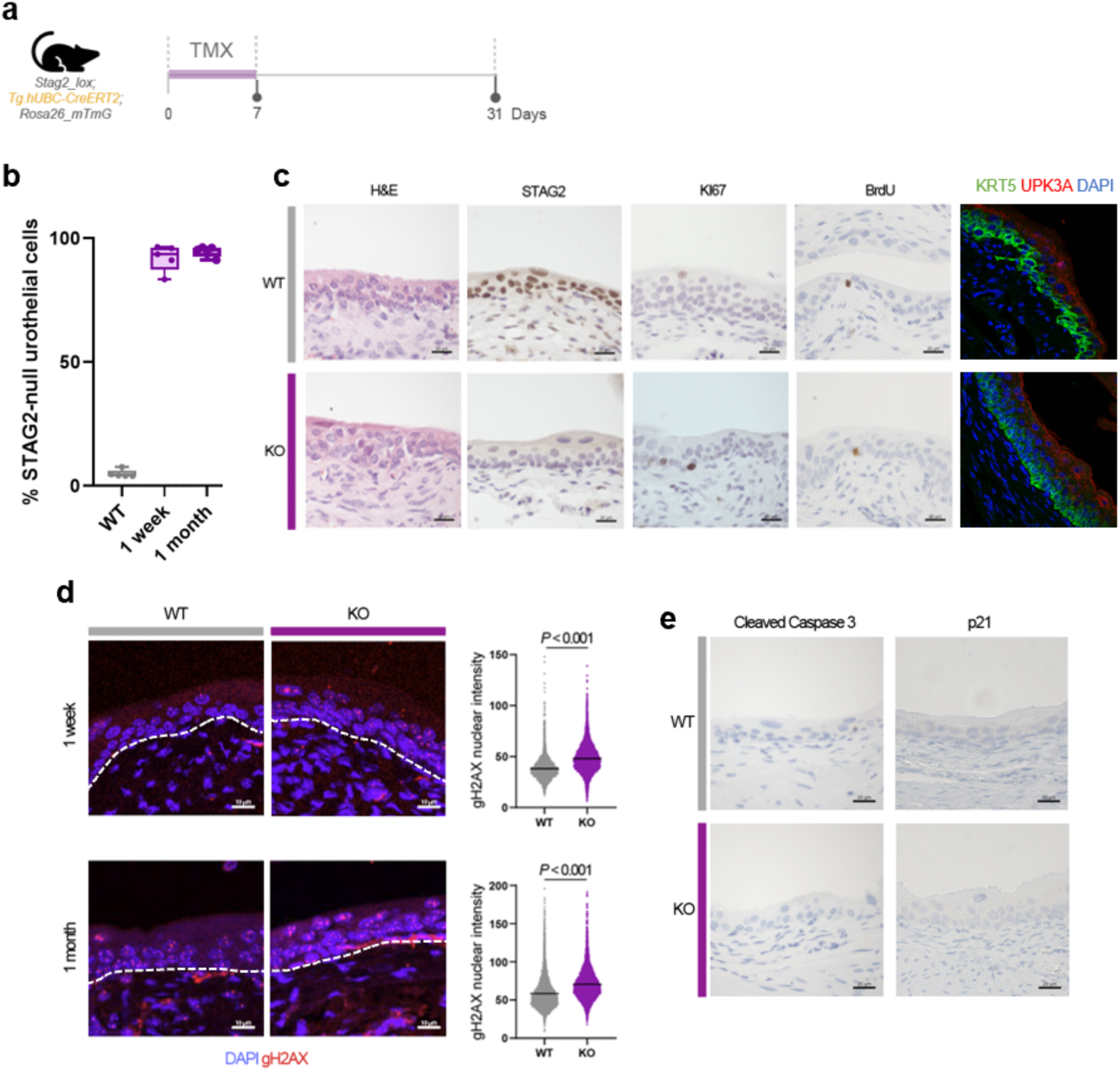
*Stag2-*KO urothelium shows increased and sustained DNA damage response with no major differences in proliferation or differentiation one month after inducing recombination. **a**, 8-12 week-old *Tg.hUBC-CreERT2;Stag2^lox/lox^;Rosa^mTmG^*mice received TMX for one week and the urothelium was analyzed at one week or one month. **b**, Box plot showing a significant increase in the percentage of STAG2-null urothelial cells in *Stag2-*KO bladder sections one week and one month after initiation of TMX administration. Each data point represents an individual mouse. **c,** H&E, IHC for STAG2, KI67, BrdU, and p63, and immunofluorescence for KRT5 and UPK3A in WT and *Stag2-*KO bladder sections at one month. Scale bar, 20 μm. **d,** immunofluorescence staining of gH2AX in WT and *Stag2*-KO bladder sections one week and one month after inducing recombination. Nuclear intensity values of gH2AX per urothelial cell and the mean per condition are plotted on the right (n = 4 mice/group). Statistical test: two-tailed Mann-Whitney. **e,** IHC analysis of cleaved caspase 3 and p21 in WT and *Stag2*-KO bladder sections, one week after inducing recombination. Scale bar, 20 μm.

### STAG2 is required for lineage stability, quiescence, and cell cycle gene repression

To identify the molecular effects of *Stag2* inactivation, we profiled the transcriptome of peeled urothelium from *Stag2 WT* and *KO* mice one week and one month after TMX administration. Despite normal histology at one week, 655 and 231 genes were significantly up-regulated and down-regulated in *Stag2*-KO urothelium, respectively (**Fig. 2a, Suppl. Table 1**). We found a significant down-regulation of a luminal/urothelial signature and a positive enrichment of the signature from Ba/Sq tumours ^29^ in *Stag2*-KO urothelium at one week and one month (**Fig. 2b**). At one week, up-regulated genes were enriched in pathways related to cell cycle (e.g., *Cdk1*, *Mki67*, *Ccnb1*, *Mcm3*, *Mcm5*, *Mcm7*, *Mcm10*) and extracellular matrix organization [e.g., collagens (e.g., *Col4a1*, *Col4a2*, *Col12a1*) and extracellular proteases (e.g., *Adamts4*, *Adamts12*, *Adam23*)] (**Fig. 2c**). Down-regulated genes were enriched in circadian rhythm-related pathways (e.g., *Per2, Dbp, Bhlhe41*) (**Fig. 2c**). Transcriptome profiling at 1 month revealed a significant up-regulation of genes related to integrin signaling (e.g., *Rras, Akt1*), focal adhesions (e.g., *Pdgfrb, Lama2*), and protein translation and metabolism (e.g., *Eif5a, Rps2*) (**Suppl. Fig. 2a,b, Suppl. Table 1**). However, there were no significant differences in expression of cell cycle genes nor pathways (**Suppl. Fig. 2a,b**), consistent with the lack of changes in Ki67 expression (**Fig. 1b, Suppl. Fig. 1c**).

**Figure 2.**
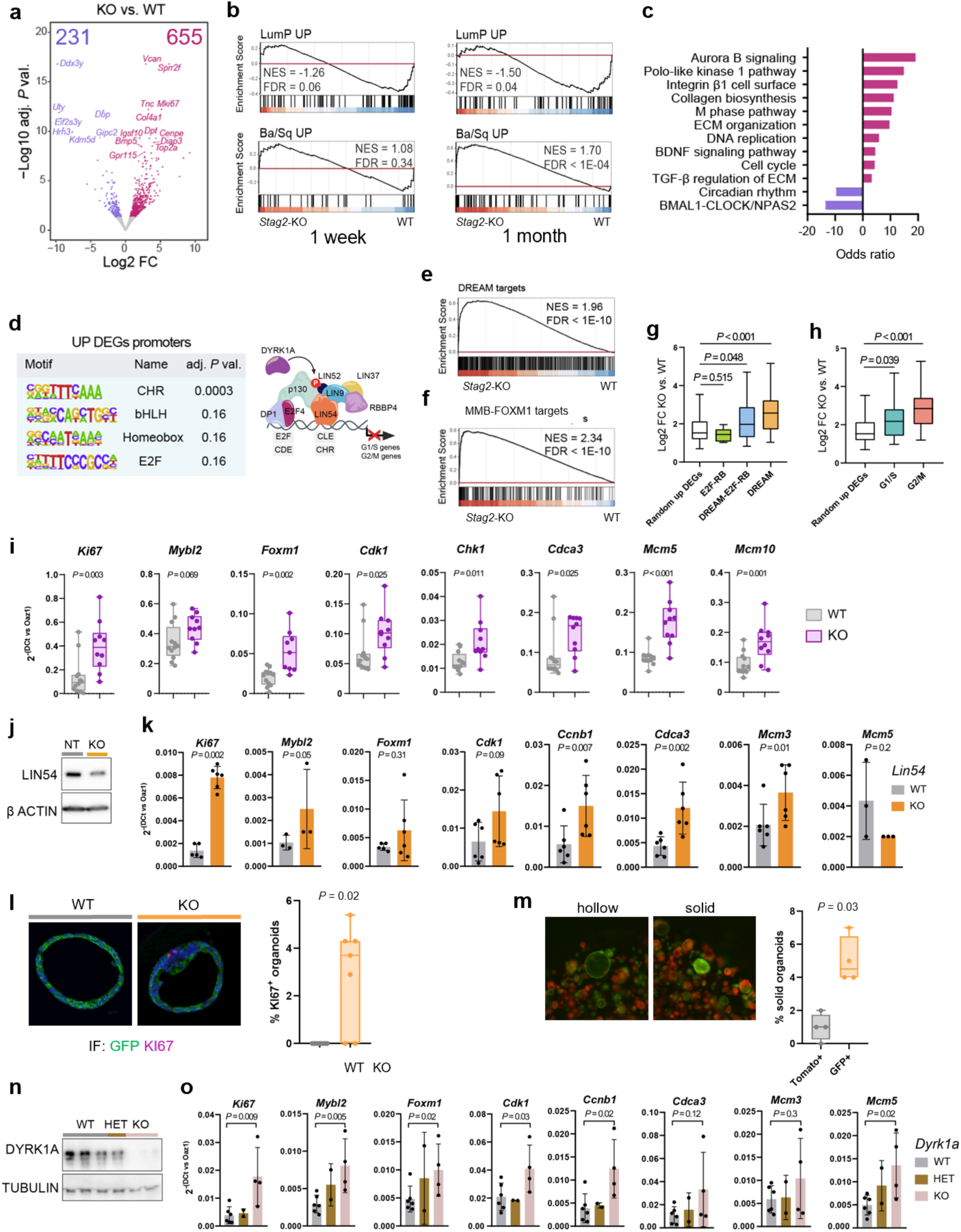
STAG2 supports urothelial lineage identity and is required for DREAM-mediated cell cycle repression and quiescence. **a-h,** 8-12 week-old *Tg.hUBC^CreERT^*^2^*;Stag2^lox/lox^;Rosa^mTmG^* mice received TMX for one week and the urothelium was analyzed at one week. **a**, Volcano plot showing gene expression changes identified in *Stag2*-KO vs. WT urothelium (adj. *P* val. < 0.05) from *Tg.hUBC^CreERT^*^2^*;Stag2^lox/lox^;Rosa^mTmG^* mice one week after inducing recombination. **b**, GSEA plots showing enrichment of the LumP and Ba/Sq up-regulated transcriptional signatures (BLCA consensus classification ^29^ in *Stag2*-KO urothelium one week (left) and one month (right) after inducing recombination. **c,** Pathway analysis showing odds ratio for the top 10 significantly up- and down-regulated pathways (adj. *P* val. < 0.05). Note that only two pathways show negative enrichment. **d**, Promoter motif analysis of up-regulated genes, showing the most significantly enriched motifs (left). Scheme depicting the DREAM complex binding the CHR motif (right). **e,f**, GSEA plots showing the up-regulation of DREAM ^30^ (**e**) and MuvB-MYB-B-FOXM1 target genes ^30^ (**f**) in *Stag2*-KO urothelium. **g**, Log2 FC expression between *Stag2*-KO vs. WT urothelium of E2F-RB (n = 5), DREAM-E2F-RB common (n = 45), and DREAM (n = 59) target genes; a set of random up-regulated genes is shown for comparison (n = 59). Statistical test: two-tailed Mann-Whitney. **h**, Log2 FC expression between *Stag2*-KO vs. WT urothelium of G1/S-phase (n = 20) and G2/M-phase (n = 50) genes; a set of random up-regulated genes is shown for comparison (n = 50). Statistical test: two-tailed Mann-Whitney. **i**, RT-qPCR analysis of expression of cell cycle genes in peeled urothelium from WT and *Stag2*-KO bladders from *Trp63^CreERT^*^2^;*Stag2^lox/lox^* mice one week after inducing recombination. Data were normalized to *Oaz1* expression. Each dot represents one biological replicate. Statistical test: two-tailed Mann-Whitney. **j**, Western blot showing expression of LIN54 five days after inducing Cas9 expression. **k,** RT-qPCR expression analysis of cell cycle genes in *Lin54*-WT and *LIN54*-KO quiescent organoids. Data were normalized to *Oaz1* expression. Each dot represents one biological replicate. Statistical test: one-tailed Mann-Whitney. **l**, KI67 expression in *Lin54*-WT and *Lin54*-KO organoids. Each dot represents one biological replicate. Statistical test: two-tailed Mann-Whitney. **m,** Percentage of solid (undifferentiated) organoids among *Lin54*-WT (Tomato^+^) and *Lin54*-KO (GFP^+^) organoids. Each dot represents one biological replicate. **n,** Western blot showing expression of DYRK1A five days after inducing recombination. **o,** RT-qPCR expression analysis of cell cycle genes in *Dyrk1a*^+/+^ (WT), *Dyrk1a^lox^*^/+^ (HET), and *Dyrk1a^lox/lox^* (KO) quiescent organoids. Data were normalized to *Oaz1* expression. Each dot represents one biological replicate. Statistical test: one-tailed Mann-Whitney.

To identify upstream cell cycle regulators, we performed promoter motif analysis of genes up-regulated at one week using HOMER. The highest enrichment corresponded to the CHR motif recognized by the DREAM complex (*adj. P* = 0.0003), followed by bHLH, homeobox, and E2F motifs (*adj. P* = 0.16) (**Fig. 2d**). GSEA revealed a highly significant up-regulation of DREAM (**Fig. 2e**) and DREAM-related activator complex MuvB-MYB-B-FOXM1 target genes (**Fig. 2f**) ^30^ in *Stag2*-KO urothelium. Up-regulation of DREAM targets was significantly higher than that of E2F-RB targets (**Fig. 2g**), in agreement with the more significant overexpression of G2/M-phase genes compared to G1/S-phase genes and the established role of MuvB-MYB-B-FOXM1 in the control of late cell cycle genes (**Fig. 2h**). Similar results were obtained using *Trp63 ^CreERT^*^2^*; Stag2^lox/lox^;Rosa26^mT/mG^* mice, indicating that *Stag2* deletion in basal and intermediate urothelial cells recapitulates the effects observed upon ubiquitous *Stag2* inactivation (**Suppl. Fig. 3a-d, Fig. 2i**), including the transient nature of changes in cell cycle gene expression (**Suppl. Fig. 4**).

To further elucidate the mechanisms involved, we established novel normal urothelial cell culture models including a spontaneously immortalized epithelial line derived from the bladder (NU1) and urothelial organoids. NU1 cells can be cultured under proliferative conditions and be induced to enter robust quiescence upon EGFR inhibition and PPARg activation, highlighting features of urothelial differentiation (**Suppl. Fig 5e-g**). The main *in vivo* findings were largely recapitulated in both *in vitro* systems.

To assess whether the transcriptomic findings reflect the effects of STAG2 loss on epithelial cells, we used urothelial organoids from *Tg.UBC^CreERT^*^2^*;Stag2^lox/lox^;Rosa26^mT/mG^* mice (**Suppl. Fig. 5a**). These organoids recapitulate key aspects of the 3D organization and function of the urothelium ^25^ and allow for synchronized cell cycle arrest upon growth factor depletion (**Suppl. Fig. 5b**) and *Stag2* inactivation using TMX (**Suppl. Fig. 5c**). Consistent with the *in vivo* findings, STAG2 loss in quiescent organoids led to the up-regulation of cell cycle genes (**Suppl. Fig. 5d**).

DREAM binds DNA through LIN54 and assembly of the complex depends on DYRK1A, a tyrosine kinase that phosphorylates Ser28 on LIN52 and allows the recruitment of hypophosphorylated p130 ^20,21,31^. To directly test the role of DREAM, we knocked out *Lin54* in quiescent urothelial organoids from *Tg.UBC^CreERT^*^2^*;Rosa26^Cas^*^9^*;Rosa26^mT/mG^* mice ^32^ using TMX (**Fig. 2j**). As with *Stag2* (**Suppl. Fig. 5d**), *Lin54* knockout resulted in increased cell cycle gene expression (**Fig. 2k**) increased numbers of KI67^+^ cells (**Fig. 2l**), and solid, undifferentiated organoids (**Fig. 2m**). *Dyrk1a* inactivation in quiescent urothelial organoids from *Tg.UBC^CreERT^*^2^*; Dyrk1a^lox/lox^* mice ^33^ also resulted in up-regulated cell cycle gene expression (**Fig. 2n,o**). These findings underscore the role of DREAM in sustaining urothelial quiescence.

**Suppl. Figure 2.**
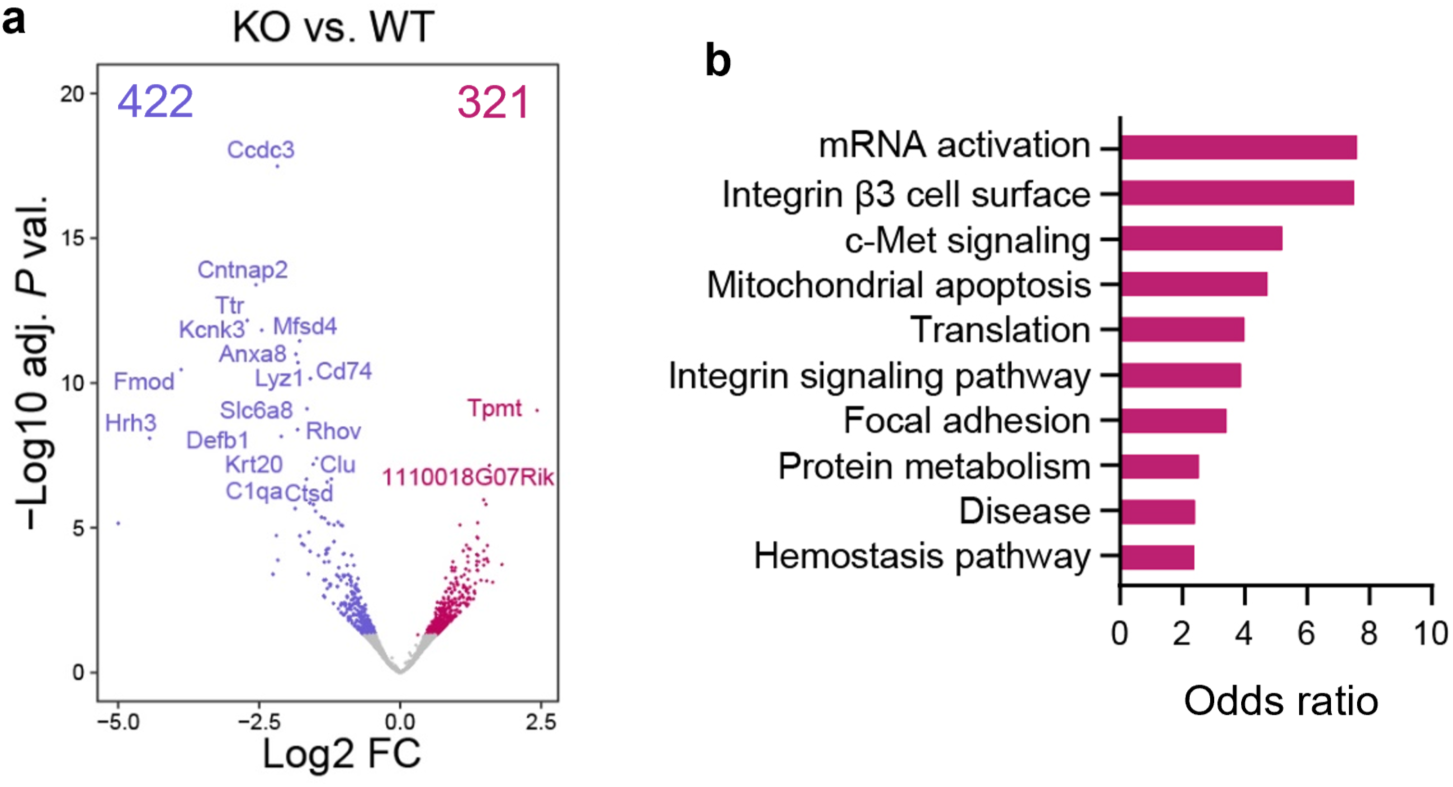
Transcriptome analysis of the urothelium one month after inducing *Stag2* recombination highlights the transient changes in expression of cell cycle genes. **a**, Volcano plot showing gene expression changes identified in *Stag2*-KO vs. WT urothelium (adj. *P* val. < 0.05) from *Tg.UBC^CreERT^*^2^*;Stag2^lox/lox^;Rosa26^mT/mG^* mice one month after inducing recombination. **b**, Pathway enrichment analysis showing odds ratio for the top 10 significantly up-regulated pathways (adj. *P* val. < 0.05).

**Suppl. Figure 3.**
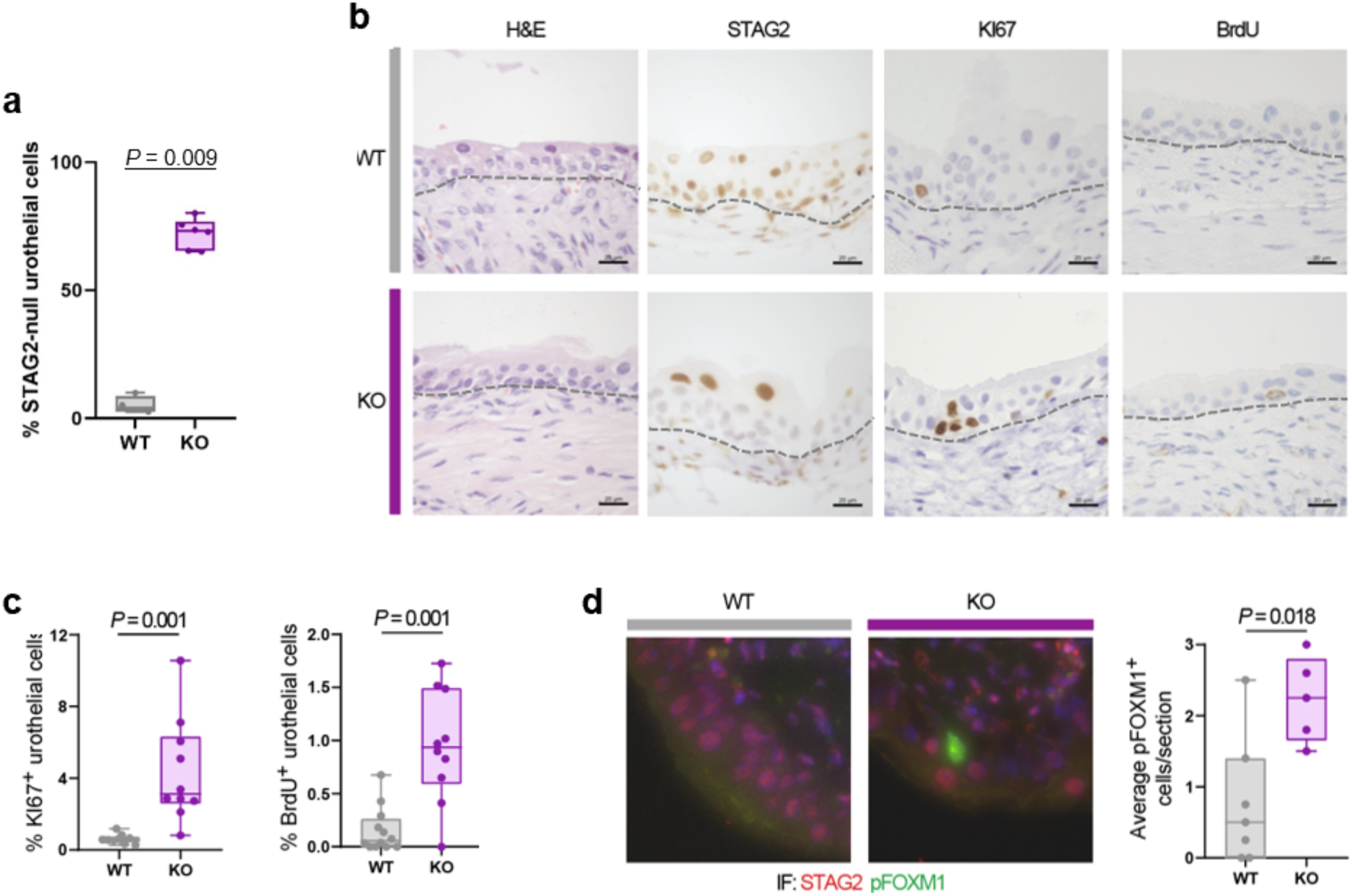
*Trp63*-driven urothelial-specific *Stag2* inactivation results in cell cycle entry without major histological changes. 8-12 week-old *Trp63 ^CreERT^*^2^*; Stag2^lox/lox^* mice received TMX for one week and the urothelium was analyzed at one week. **a**, Box plot showing a significant reduction in STAG2^+^ urothelial cells in *Stag2-*KO bladder sections. Each data point represents an individual mouse. Statistical test: two-tailed Mann-Whitney. **b**, H&E and IHC analysis of STAG2, KI67, and BrdU in WT and *Stag2-*KO bladder sections. Scale bar, 20 μm. **c**, Percentage of KI67^+^ and BrdU^+^ urothelial cells in WT and *Stag2-*KO bladder sections. Each data point represents an individual mouse. Statistical test: two-tailed Mann-Whitney. **d**, IF analysis of STAG2 (red) and pFOXM1 (green) and quantification of pFOXM1^+^ cells in bladder sections. Each data point represents an individual mouse. Statistical test: two-tailed Mann-Whitney.

**Suppl. Fig 4.**
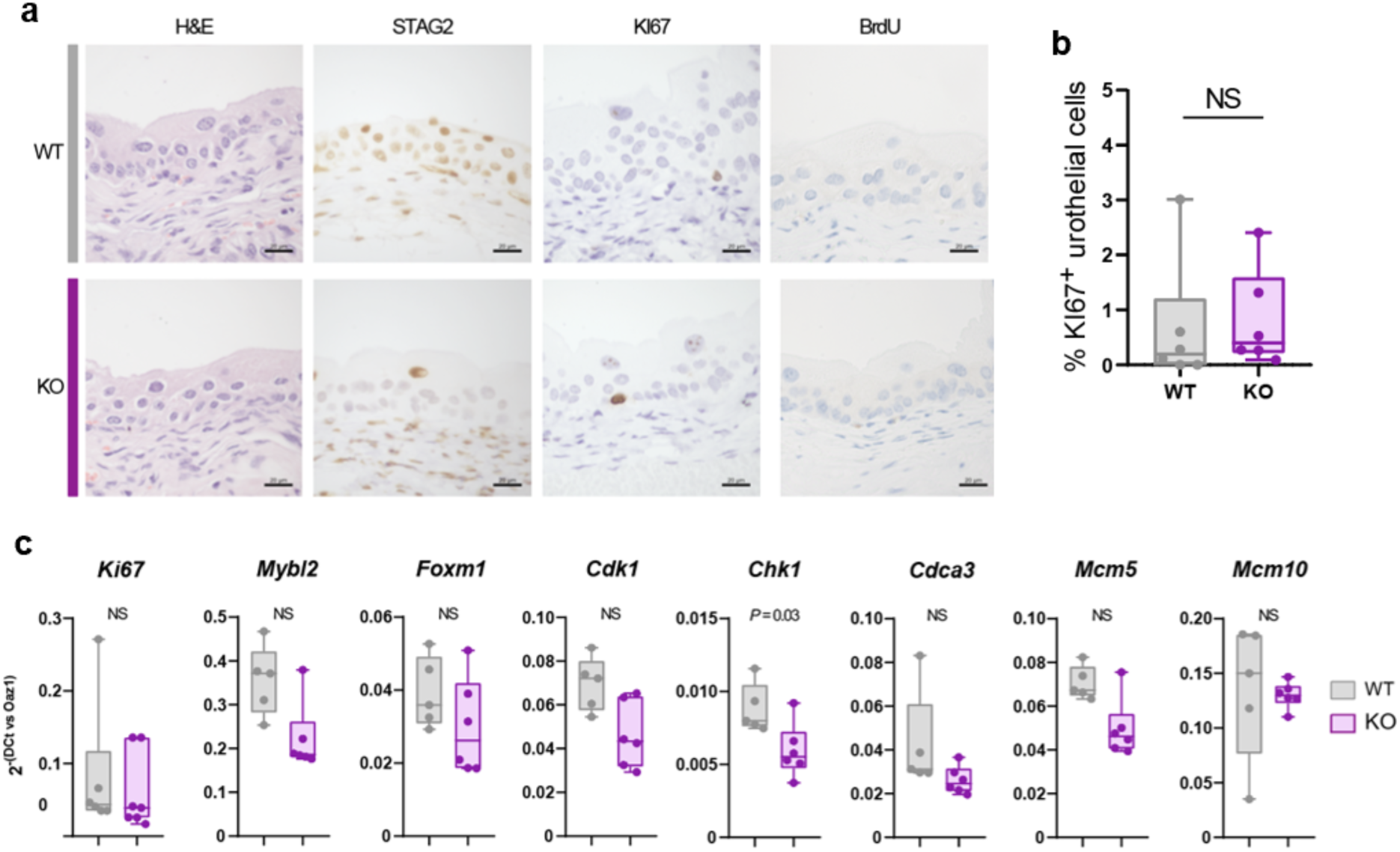
Cell cycle gene expression returns to basal levels one month after urothelial-restricted *Stag2* inactivation. 8-12 week-old *Trp63 ^CreERT^*^2^*;Stag2^lox/lox^* mice received TMX for one week and the urothelium was analyzed at one month. **a**, H&E and IHC analysis of STAG2, KI67, and BrdU in WT and *Stag2-*KO bladder sections. Scale bar, 20 μm. **b**, Percentage of KI67^+^ urothelial cells in WT and *Stag2-*KO bladder sections. Each data point represents an individual mouse. Statistical test: two-tailed Mann-Whitney. **c**, RT-qPCR analysis of expression of cell cycle genes in peeled urothelium. Data were normalized to *Oaz1* expression. Each dot represents one biological replicate. Statistical test: two-tailed Mann-Whitney.

**Suppl. Figure 5.**
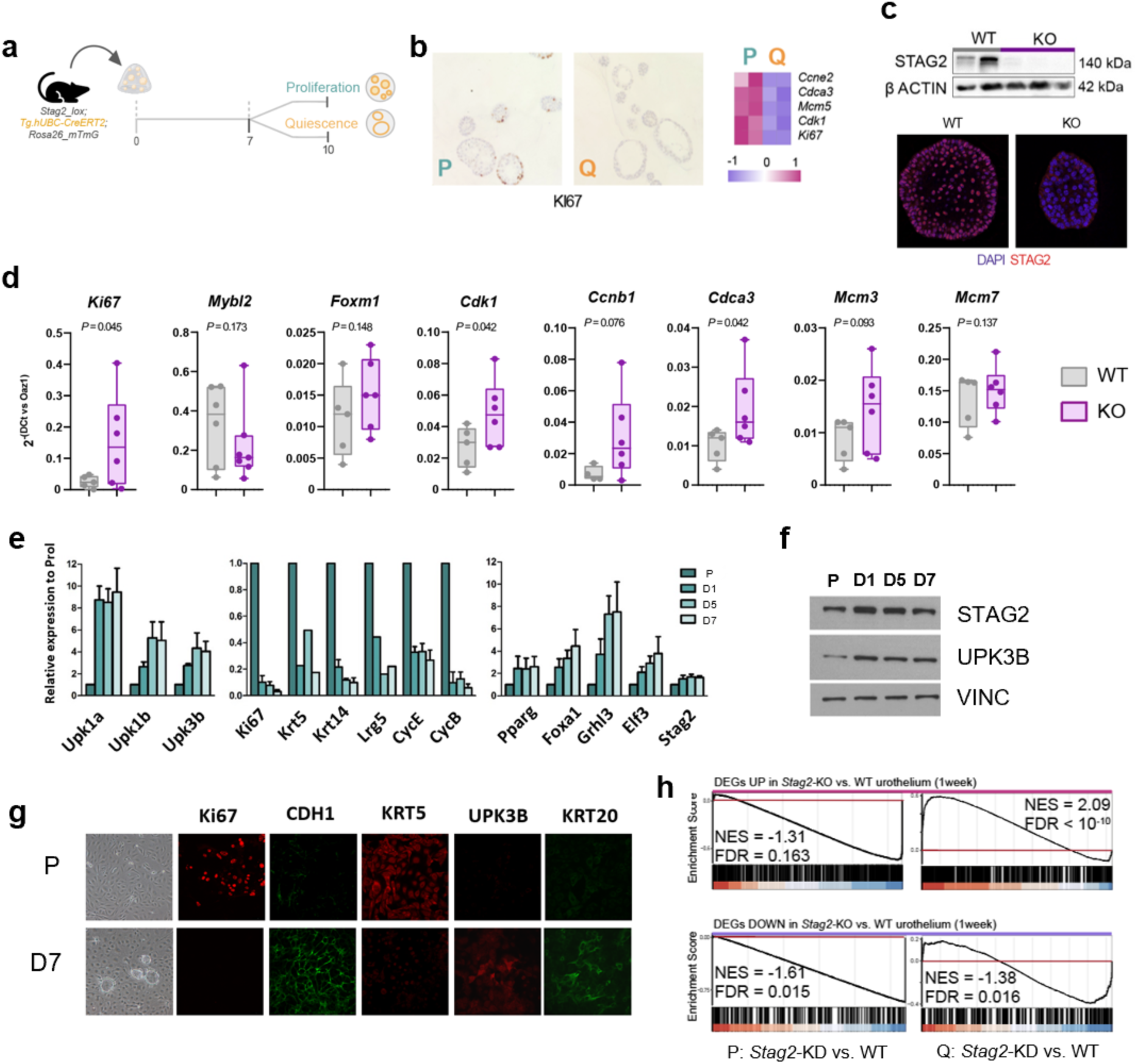
Validation of urothelial organoids and NU1 cells as model systems to assess the role of STAG2 in urothelial biology. **a-d,** Normal mouse urothelial organoids. **a,** Experimental design: urothelial cells from mouse bladders were cultured as organoids as described ^25^ and maintained in either complete medium (proliferation; P) or in growth factor-depleted medium (quiescence; Q) for 3 days. **b,** Cell cycle marker expression in proliferative and quiescent organoids. **c**, Lack of expression of STAG2 in organoids upon induction of recombination with TMX. **d**, RT-qPCR analysis of expression of cell cycle genes in *Stag2*-KO and WT quiescent urothelial organoids. Each data point represents one biological replicate. Statistical test: one-tailed Mann-Whitney. **e-h**, NU1 cells. **e**, Expression of urothelial differentiation markers in spontaneously immortalized NU1 cells after 1, 5, or 7 days of induction of differentiation (D); expression relative to proliferating (P) cells (n=3). **f,** Western blot analysis of STAG2 and UPK3B expression in NU1 cells in proliferation (P) and 1, 5, and 7 days after induction of quiescence/differentiation (D). **g,** IF analysis of KI67, CDH1, KRT5, UPK3B, and KRT20 in proliferating and differentiated NU1 cells. **h,** GSEA plots showing the enrichment of signatures of genes up- and down-regulated in *Stag2*-KO urothelium at 1 week in *Stag2*-KD proliferating (P; left) and quiescent (Q; right) NU1 cells.

### STAG2 is selectively enriched at promoters of cell cycle genes

To elucidate the genomic mechanisms involved, we used the spontaneously immortalized murine NU1 cells. *Stag2* was effectively knocked down (KD) in NU1 cells using lentiviral shRNAs (**Fig. 3**), in proliferative or quiescent conditions and the transcriptome was analyzed (**Fig. 3a, b**). Only 20 genes were differentially expressed in *Stag2*-KD vs. sh-control proliferative cells (FDR < 0.05) (**Fig. 3a, Suppl. Dataset 1**). In contrast, 1,013 DEGs were differentially expressed in quiescent cells: 470 genes were up-regulated and 543 genes were down-regulated (**Fig. 3b, Suppl. Dataset 1**). Transcriptional changes identified *in vivo* were largely recapitulated in *Stag2*-KD quiescent cells (**Suppl. Fig. 5h**): up-regulated genes were significantly enriched in cell cycle pathways (**Fig. 3c**) and DREAM and MuvB-MYB-B-FOXM1 target genes (**Fig. 3d**), and western blotting showed increased expression of E2F1, CCNE1, MCM3, CDC6, and CHEK1 in *Stag2*-KD quiescent cells, but not in *Stag2*-KD proliferating cells (**Fig. 3e**). This indicates that *Stag2* inactivation has a much greater impact on quiescent urothelial cells.

**Figure 3.**
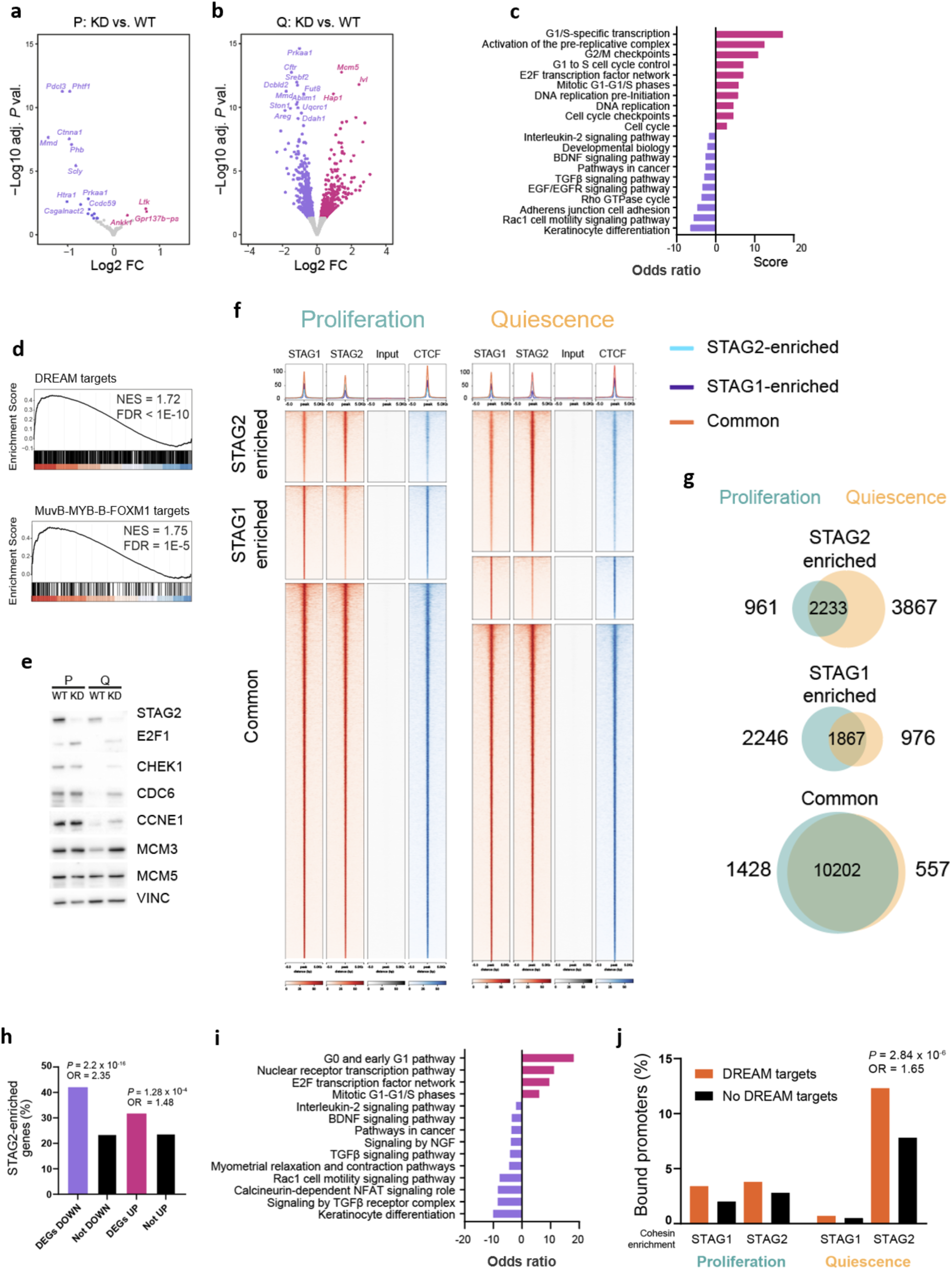
STAG2-cohesin regulates cell cycle gene programs and binds the promoter of DREAM targets in quiescent urothelial cells. **a,b**, Volcano plots showing gene expression changes identified in *Stag2*-KD vs. WT NU1 cells, in proliferation (**a**) and in quiescence after 7 days of differentiation (**b**). **c**, Pathway enrichment analysis in *Stag2*-KD vs. WT quiescent NU1 cells. **d**, GSEA plots showing the up-regulation of DREAM ^30^ (upper) and MuvB-MYB-B-FOXM1 target genes ^30^ (lower) in *Stag2*-KD cells. **e**, Western blot showing increased expression of cell cycle proteins in quiescent NU1 cells upon *Stag2* KD. VINC is included as a loading control. **f**, Clustered heatmaps displaying the genomic distribution of STAG2, STAG1, and CTCF signal (Z-score normalized) at STAG2-enriched, STAG1-enriched, and common peaks. Top plots show average signal distribution of STAG2-enriched, STAG1-enriched, and common positions (Z-score normalized). **g**, Venn diagrams showing the overlap for STAG2-enriched, STAG1-enriched, and common positions in proliferation and quiescence. **h**, STAG2-enriched regions are significantly over-represented among genes differentially expressed in *Stag2*-KD quiescent cells. **i**, Pathway analysis of STAG2-enriched genomic regions at DEGs (top 10 significantly up- and down-regulated pathways) (adj. *P* val. < 0.05). **j**, STAG2-enriched regions, unlike STAG1-enriched, are significantly over-represented at DREAM target gene promoters in quiescent, but not in proliferating, cells. *P* values and odds ratios (OR), Fisher’s exact test.

We then profiled the genome-wide distribution of STAG2, STAG1, and CTCF in proliferative and quiescent NU1 cells using ChIP-Seq (**Fig. 3f**). STAG2-enriched positions showed the highest distribution at promoters compared to STAG1-enriched and common positions (**Suppl. Fig. 6a**) and reduced CTCF binding (**Fig. 3f**), consistent with a role in transcriptional regulation ^34–36^. STAG2-enriched regions were significantly increased in quiescent cells, with 3867 gained binding sites (**Fig. 3g**). AP1 and SP1/KLF motifs were over-represented among STAG2-enriched positions spanning promoters and AP1 motifs were also enriched at non-promoter regions (**Suppl. Fig. 6b,c**). STAG1-enriched and common cohesin positions colocalized with CTCF in proliferative and quiescent cells (**Fig. 3f**), as described in other cell types ^37^. They were mainly found in introns and intergenic regions in both proliferative and quiescent cells (**Suppl. Fig. 6a**) and were enriched in binding motifs for CTCF and BORIS (**Suppl. Fig. 6b**).

In quiescent cells, STAG2-enriched genomic regions were over-represented among genes differentially expressed in *Stag2*-KD quiescent cells (Fisher’s exact test; *P* < 0.001, odds ratio > 1.48) (**Fig. 3h**). They were present in 34% and 44% of significantly up- and down-regulated genes, respectively (**Fig. 3h**). STAG2-enriched positions at up-regulated DEGs included cell cycle-related genes (e.g., *Mybl2, Ccnb1, Ccne2*) whereas those at down-regulated DEGs included genes related to differentiation, TGFβ signaling, and cell adhesion and motility (e.g., *Smad2, Snai2, Pik3r2*) (**Fig. 3i, Suppl. Fig. 6d**). STAG2 was significantly enriched in promoters annotated to DREAM-target genes (e.g., *Mybl2, Ccnb1*) in quiescent (odds ratio = 1.65, *P* = 2.84E-06) but not in proliferating (odds ratio = 1.39, *P* = 0.06) cells (**Fig. 3j**). These findings support a repressive role of STAG2, together with DREAM, at promoters of cell cycle genes in quiescence.

**Suppl. Figure 6.**
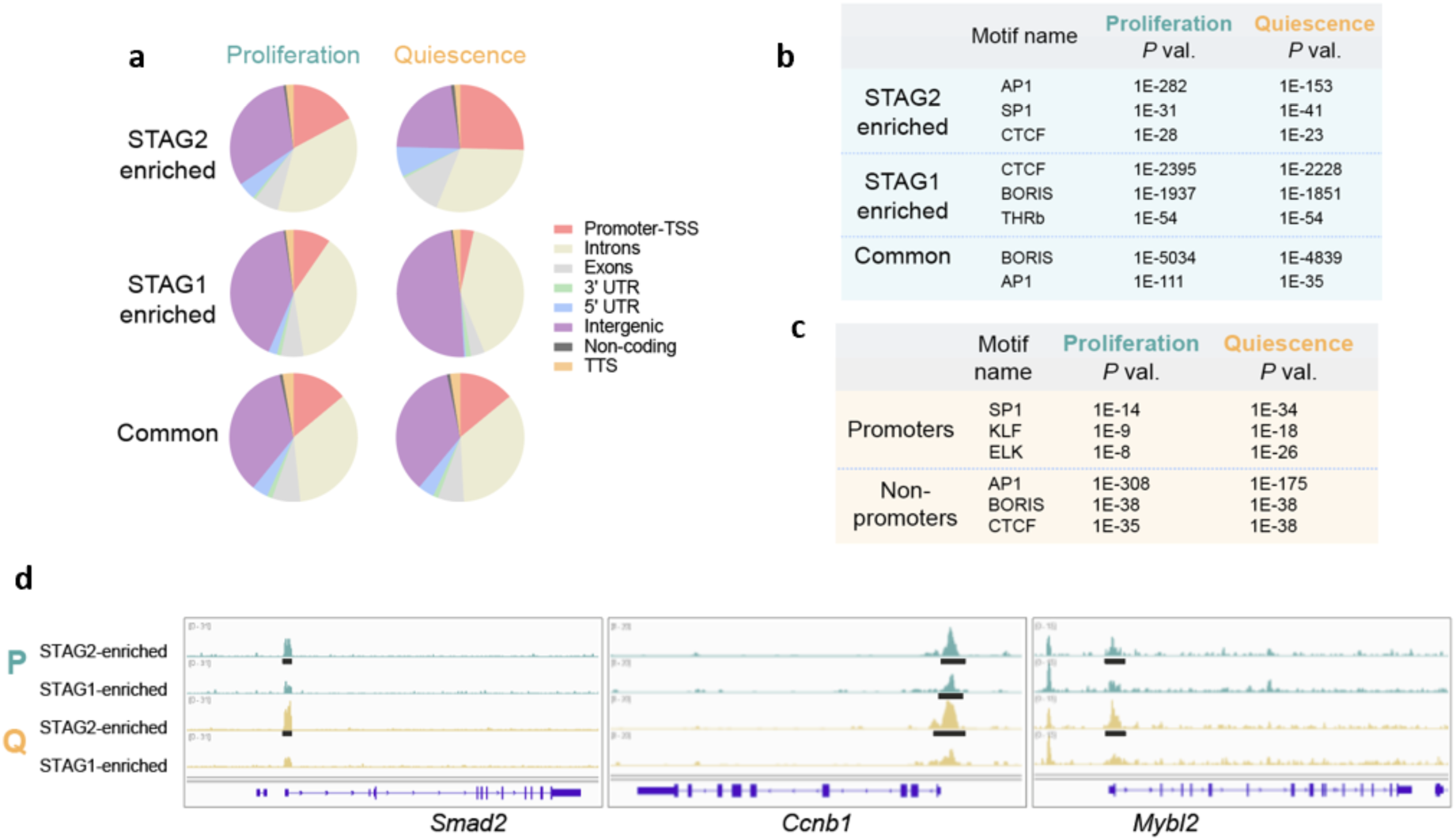
Genomic distribution of STAG2- and STAG1-cohesin in proliferating and quiescent urothelial cells reveals unique and overlapping features. **a**, Genomic annotation for STAG2-enriched, STAG1-enriched, and common positions in proliferating and quiescent NU1 cells. **b,** Motif analysis of all STAG2-enriched, STAG1-enriched, and common positions. **c**, Motif analysis of STAG2-enriched positions at promoter and non-promoter regions in proliferation and quiescence. The top 3 most significantly enriched motifs identified are shown (adj. *P* val < 0.001). **d**, Genome browser screenshot showing ChIP-Seq peak distribution for STAG2- and STAG1-enriched positions in representative genes in proliferative vs. quiescent cells. Black lines indicate significant peaks (FDR < 0.05, fold enrichment > 10, pileup > 30). One representative replicate per condition.

### STAG2-dependent programmes are conserved in normal mouse and human urothelium

To determine whether the STAG2-dependent programmes identified in mouse urothelium are conserved in human urothelium, we compared the transcriptome of normal human bladder samples (GTEx consortium) with the lowest and highest *STAG2* mRNA expression levels (n = 5/group; out of 21 samples in the GTEx database) (**Fig. 4a**). A total of 1,839 and 1,832 genes were differentially expressed at higher or lower levels in the *STAG2*-low group, respectively (**Fig. 4b**). The signatures of STAG2-dependent genes described above (**Fig. 2a**) were significantly enriched, in the corresponding direction, in *STAG2*-low vs. *STAG2*-high human samples (**Fig. 4c**). Signatures of up- and down-regulated genes in *Stag2-* KD vs. control quiescent NU1 cells, as well as DEGs bound by STAG2-cohesin (data not shown), also showed consistent up- and down-regulation in *STAG2*-low vs. *STAG2*-high samples (**Fig. 4d**). To assess whether the signatures of STAG2-dependent genes are relevant to tumour initiation, we compared the transcriptomes of non-neoplastic samples adjacent to tumour from the bladder TCGA cohort to healthy GTEx samples ^38^. We found a significant enrichment in the signatures of genes that were up- and down-regulated in *Stag2*-KO vs. WT mouse urothelium (**Fig. 4e**).

**Figure 4.**
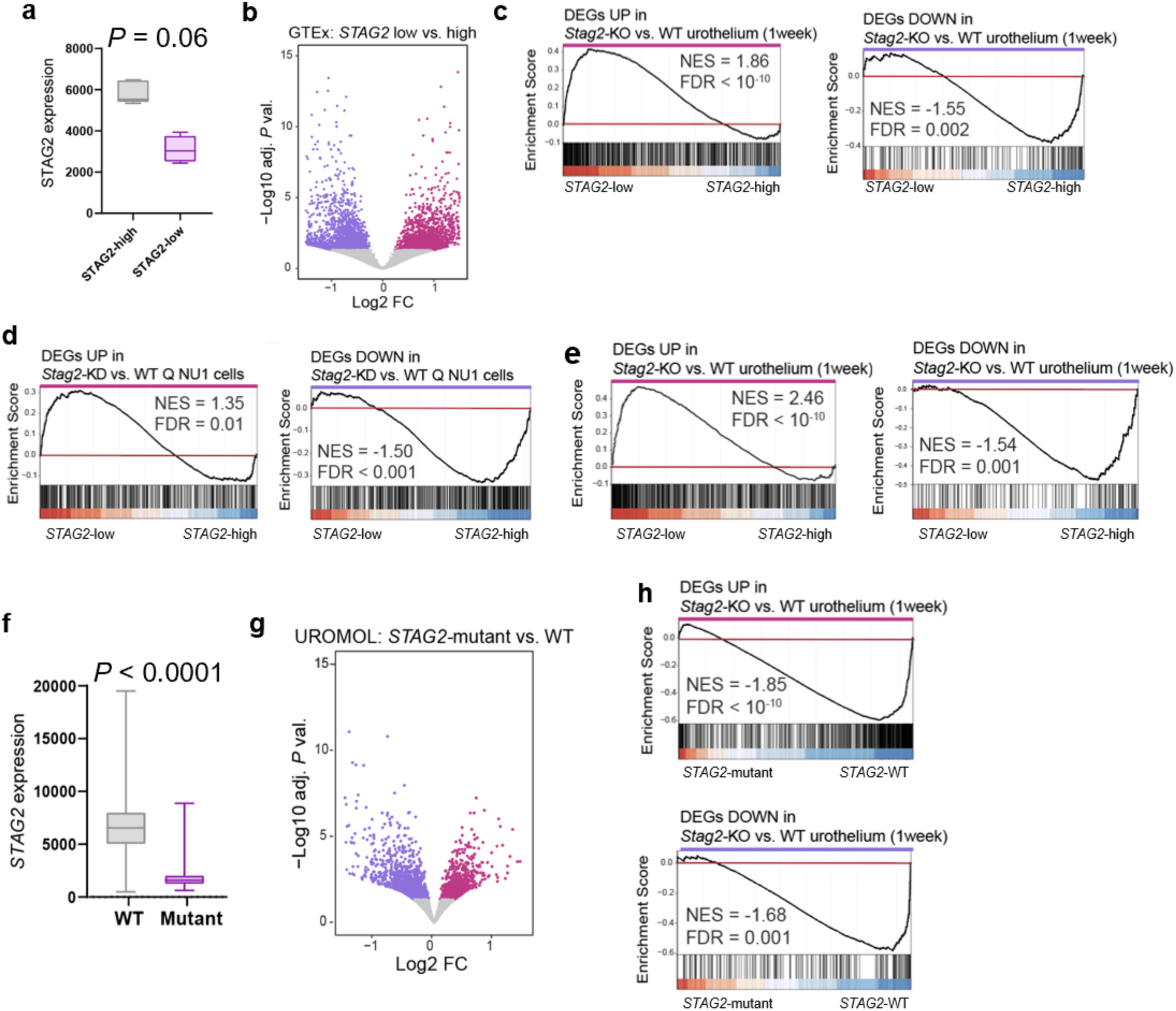
STAG2-dependent urothelial programs are conserved in normal human bladder. **a**, Expression of *STAG2* in GTEx samples with lowest and highest quartiles of *STAG2* mRNA expression (n = 5 per group). **b**, Volcano plot showing differentially expressed genes in *STAG2*-low vs. *STAG2*-high human bladders. **c,** GSEA plot showing that signatures of genes up-regulated or down-regulated in *Stag2*-KO vs. WT urothelium (one week) are positively or negatively enriched in *STAG2*-low vs. *STAG2*-high human bladders, respectively. **d,** GSEA plot showing that signatures of genes up-regulated or down-regulated in *Stag2*-KD vs. control quiescent NU1 cells are positively or negatively enriched in *STAG2*-low vs. *STAG2*-high human bladders, respectively. **e,** GSEA plot showing that signatures of genes up-regulated or down-regulated in *Stag2*-KO vs. WT urothelium (one week) are positively or negatively enriched in non-neoplastic urothelium adjacent to tumour vs. normal urothelium. **f,** *STAG2* expression levels in *STAG2-*mutant (n = 60) and WT (n = 96) low-grade tumours from the UROMOL2 study. **g,** Volcano plot showing gene expression changes identified in *STAG2*-mutant vs. WT tumours. **h,** GSEA plots showing the enrichment of DEGs found in *Stag2*-KO mouse urothelium one week after inducing recombination when comparing *STAG2*-mutant vs. WT tumours. GTEx: Genotype-Tissue Expression project, Q: quiescent.

We next assessed the STAG2-dependent gene signatures in samples from UROMOL2, the largest study of non-muscle-invasive bladder cancer. We focused on Ta low-grade tumours, the subtype with highest frequency of *STAG2* mutations ^2,3^. As expected, *STAG2* mutant samples, identified using whole exome sequencing (mutant, n = 60; WT, n = 96), displayed significantly lower levels of *STAG2* mRNA (*P* <0.0001) (**Fig. 4f**) and major transcriptomic differences (**Fig. 4g**). The signature comprising genes up-regulated in *Stag2-KO* mouse urothelium at one week was negatively enriched in *STAG2*-mutant vs. *STAG2*-WT low-grade tumours (NES = −1.85, FDR < 10-4), as was the signature of down-regulated genes (NES = −1.68, FDR = 0.001) (**Fig. 4h**). These findings support conservation of STAG2-dependent programs in normal human bladder as well as distinct roles of STAG2 in cancer cells compared to normal tissue.

### STAG2 is required for DREAM-mediated repression of cell cycle genes in quiescent cells

To dissect the molecular mechanisms underlying transcriptional changes upon *Stag2* depletion in quiescent urothelial cells *in vivo,* we performed ATAC-Seq on peeled urothelium one week post-recombination. Three hundred and 1,043 regions gained or lost accessibility, respectively, located mainly in intergenic or intronic regions (**Fig. 5a,b**). Ninety-two percent of the DEGs in *Stag2-KO* urothelium, and 96% of up-regulated DREAM target genes (data not shown) were annotated to genomic regions that were accessible in WT cells and remained so upon *Stag2* inactivation (**Fig. 5c**). This suggests that other mechanisms are responsible for activation of gene expression. We thus assessed whether the up-regulation of cell cycle genes could result from changes in DREAM genomic distribution. We performed Cut&Run for endogenous STAG2 and LIN54 using urothelial cells from WT and *Stag2-KO* bladders one week post-recombination (**Suppl. Fig. 7a**). We found a significant overlap between STAG2 and LIN54 binding (OR = 4.71, *P* < 2E-16) (**Fig. 5d,e**), 80% of LIN54-bound regions also being bound by STAG2. These co-bound regions were located at promoters (37%), introns (33%), and intergenic regions (21%) (**Fig. 5e,f**). STAG2-LIN54 shared positions mapped to 9.6% of genes up-regulated in *Stag2*-KO urothelium and were enriched in promoters of cell cycle genes (**Fig. 5g,h**).

**Figure 5.**
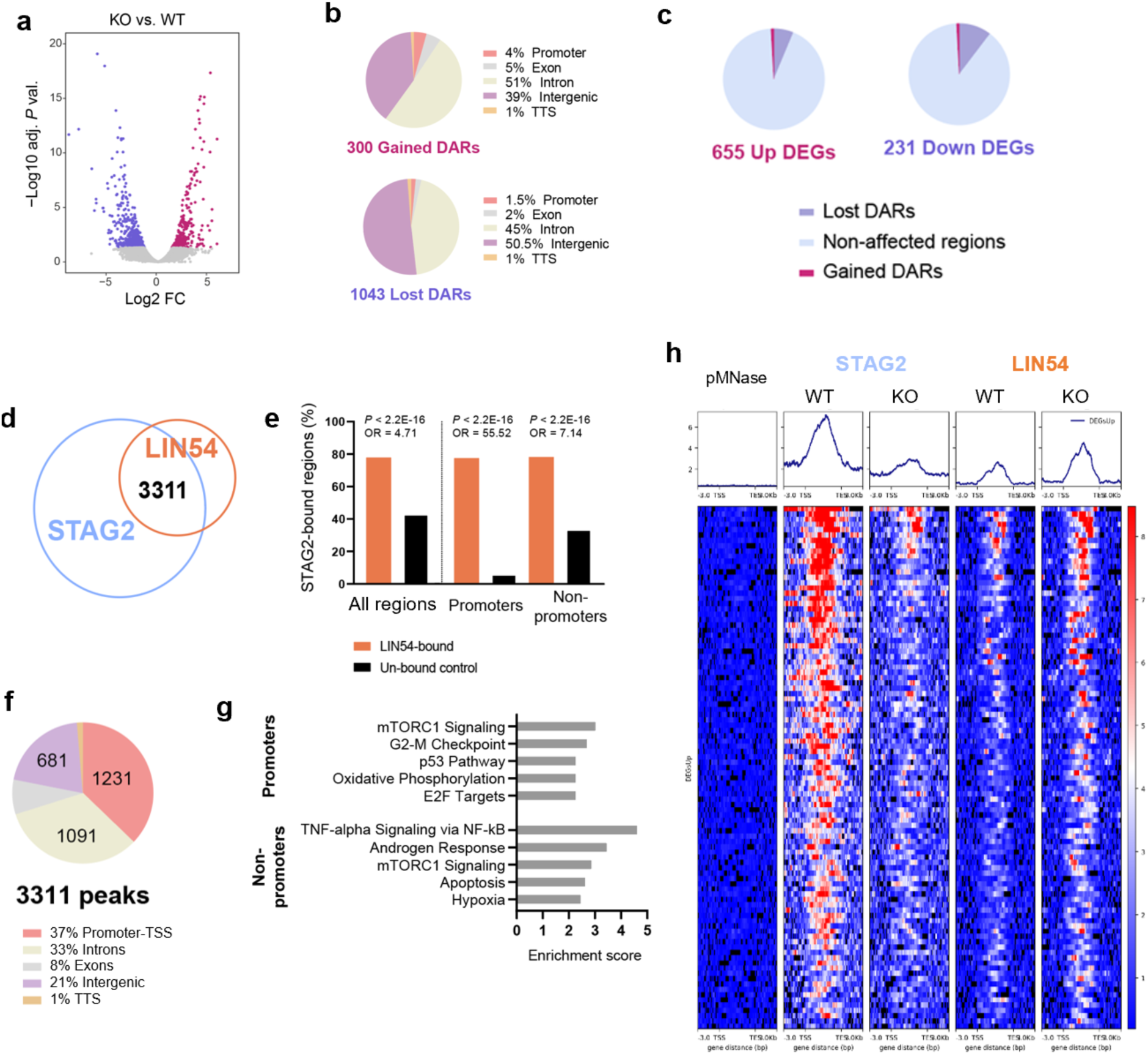
STAG2 and DREAM share genomic binding to the promoter of cell cycle genes. **a**, Volcano plot showing changes in chromatin accessibility in *Stag2*-KO urothelium (adj. P val. < 0.05). **b**, Genomic annotation for gained and lost DARs. **c,** Changes in gene expression do not associate closely with changes in chromatin accessibility. **d**, Overlap between consensus peaks for STAG2 and LIN54 in WT urothelial cells. **e**, STAG2-bound promoter and non-promoter regions occupied by LIN54. **f**, Genomic annotation for STAG2-DREAM shared sites. **g,** Top five significantly over-represented pathways among genes annotated to STAG2-DREAM shared sites (adj. P val. < 0.05) based on the Hallmarks collection (MSigDB, 2020). **h**, Heatmap depicting pMNase, STAG2, and LIN54 signals (Z-score normalized) in WT and *Stag2*-KO samples at STAG2-bound regions annotated to genes up-regulated in *Stag2*-KO urothelium. Top panel, average signal plots are shown.

Differential binding analysis revealed 201 and 3 regions with significantly increased or decreased LIN54 occupancy in *Stag2-KO* vs. WT cells (FDR < 0.05), respectively (**Suppl. Fig. 7b-d**), mostly spanning promoters, intergenic regions, and introns (**Suppl. Fig. 7e**). We next compared LIN54 peaks at up-regulated DEGs in *Stag2*-KO vs. WT cells (**Fig. 6a**) and identified 101 sites with increased LIN54 signal (FDR < 0.1; none at FDR < 0.05) (**Fig. 6b**), preferentially spanning introns (44.5%), promoters (26%), and intergenic regions (27%) (**Fig. 6c**). Differential peaks annotated to promoters were enriched in cell cycle genes (e.g., *Mcm10, Ndc80, Nas*p, and *Ncapg2*) and pathways, as well as in KLF, SP, and E2F1 motifs (**Fig. 6d-f**). In contrast, those annotated to non-promoter regions were enriched in AP1 motifs (**Fig. 6e**). Interestingly, AP1 motifs have also been reported to be enriched in Chip-Seq peaks for RB at non-promoter sites in cycling cells ^39^. Altogether, these data indicate that *Stag2* inactivation results in altered LIN54 distribution and increased genome-wide occupancy, with a modest gain in LIN54 binding at promoters of cell cycle genes that are up-regulated upon STAG2 loss. To assess whether STAG2 loss results in changes in DREAM complex composition, we performed immunoprecipitation followed by mass spectrometry (IP-MS) for endogenous LIN9, the DREAM scaffolding protein ^40^, in WT and *Stag2*-KO quiescent urothelial organoids. We found no evidence of major changes in expression levels of core DREAM complex components at the mRNA or protein level (**Suppl. Fig. 8a,b**). However, we found a reproducibly reduced interaction between LIN9 and LIN52 (P < 0.05) in three independent experiments, as well as a lower interaction with p130 (**Fig. 6g, Suppl. Fig. 8c, Suppl. Dataset 2**). Considering that LIN52 is crucial for p130 recruitment, DREAM assembly, and cell cycle gene repression ^20,21^ the reduced LIN52 (and p130) binding in the absence of STAG2 may contribute to the de-repression of cell cycle genes. STAG2 was not found among LIN9 interactors in our experiments (**Suppl. Dataset 2**) nor did it co-immunoprecipitate with LIN54 in RT112 bladder cancer cells, which express WT STAG2 (**Fig. 6h**). This suggests that other mechanisms are involved in impaired DREAM function upon STAG2 loss, including changes in chromatin organization.

**Figure 6.**
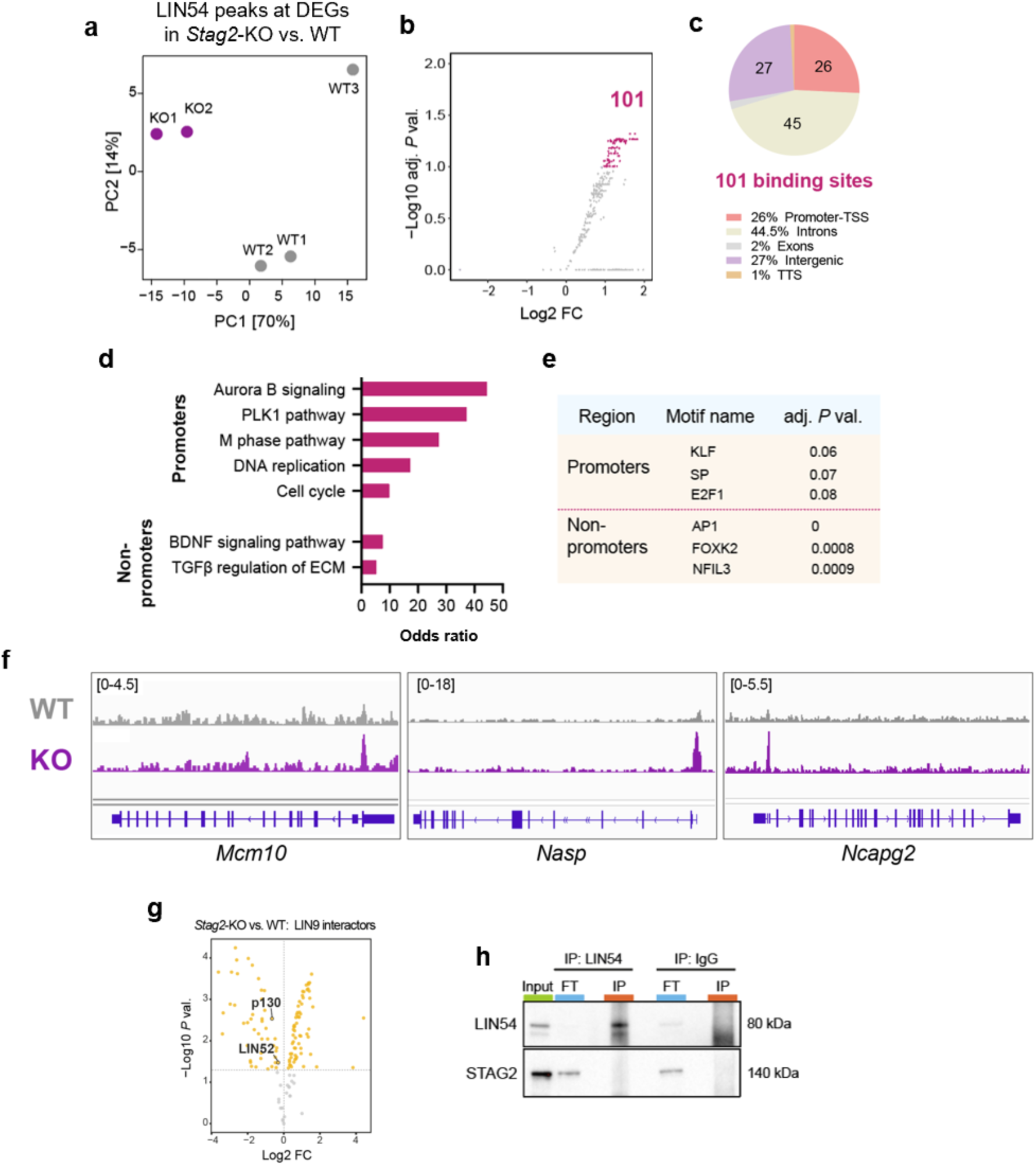
*Stag2* inactivation results in altered DREAM genomic distribution and composition. **a,** Principal component analysis restricted to regions annotated to DEGs up-regulated in *Stag2*-KO urothelium one week upon inducing recombination. **b**, Volcano plot showing changes in LIN54 binding annotated to genes up-regulated in *Stag2*-KO vs. WT urothelium (FDR < 0.1). **c**, Genomic annotation of regions with increased LIN54 binding from panel (b). **d**, Pathway analysis showing odds ratio for the top significantly over-represented pathways among up-regulated DEGs annotated to regions with increased LIN54 binding (adj. *P* val. < 0.05). **e,** Motif enrichment analysis of regions with increased LIN54 binding. The top 3 most enriched motifs identified are shown. **f**, Genome browser screenshot showing LIN54 binding signal to exemplary regions in WT and *Stag2*-KO cells. One representative replicate per condition is shown. **g,** Volcano plot showing differential LIN9 interactors in *Stag2*-KO vs. WT quiescent organoids identified by IP-MS analysis. LIN52 is highlighted (P val. < 0.05). Results from a representative experiment. **h**, Western blot showing the results of co-immunoprecipitation analysis to assess the interaction of LIN54 and STAG2 in RT112 bladder cancer cell lysates. IP: immunoprecipitation; FT: flow through.

**Suppl. Fig 7.**
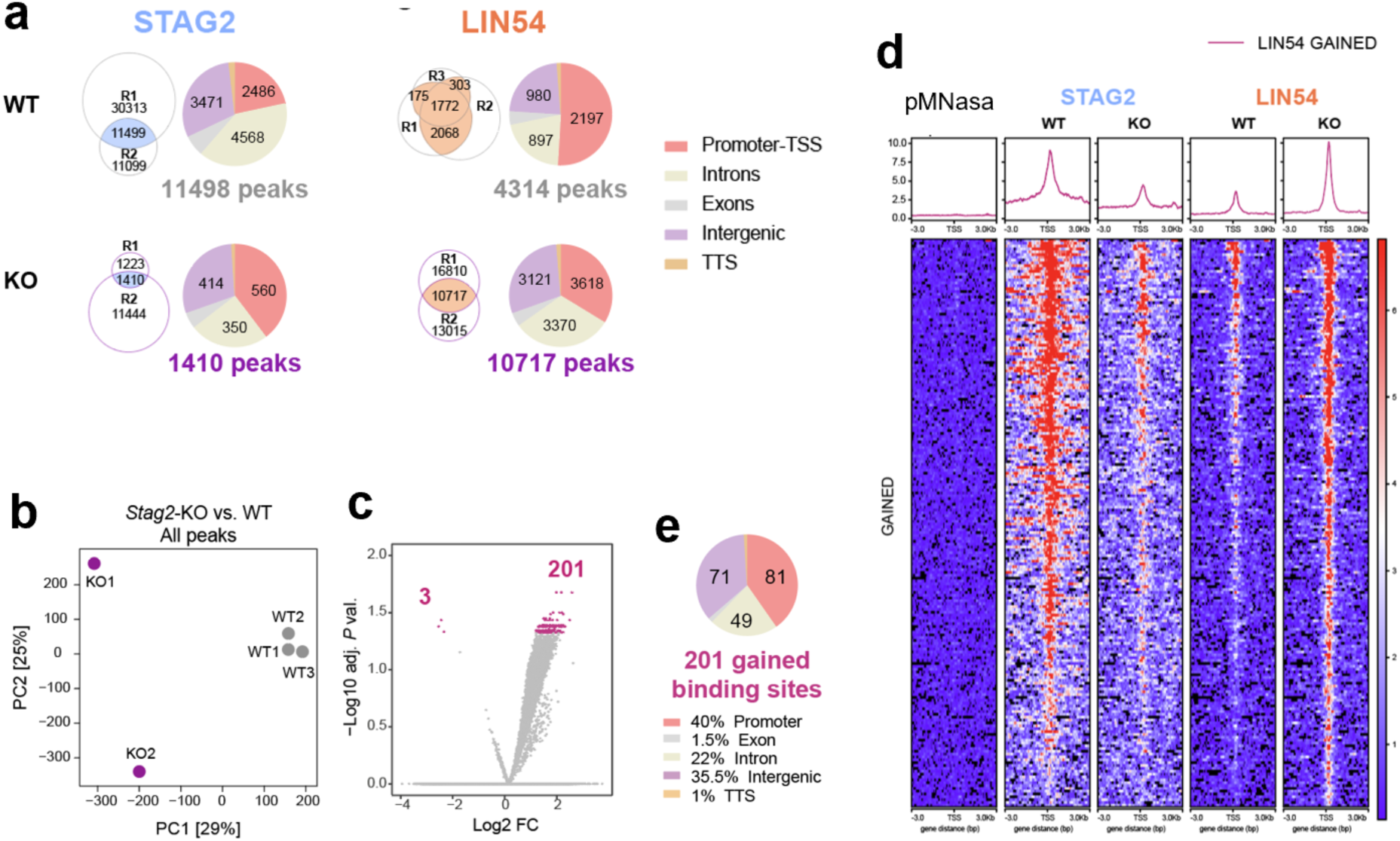
STAG2 loss results in increased LIN54 genome wide binding. **a,** Venn diagrams depicting the genomic annotation of STAG2 and LIN54 binding regions in WT and *Stag2*-KO cells. **b,** Principal component analysis. **c,** Volcano plot showing changes in LIN54 binding identified in *Stag2*-KO vs. WT urothelium (FDR < 0.05). **d**) Heatmap showing LIN54 signal at gained sites **e**, Genomic annotation of regions with increased LIN54 binding upon STAG2 loss (FDR < 0.05).

**Suppl. Figure 8.**
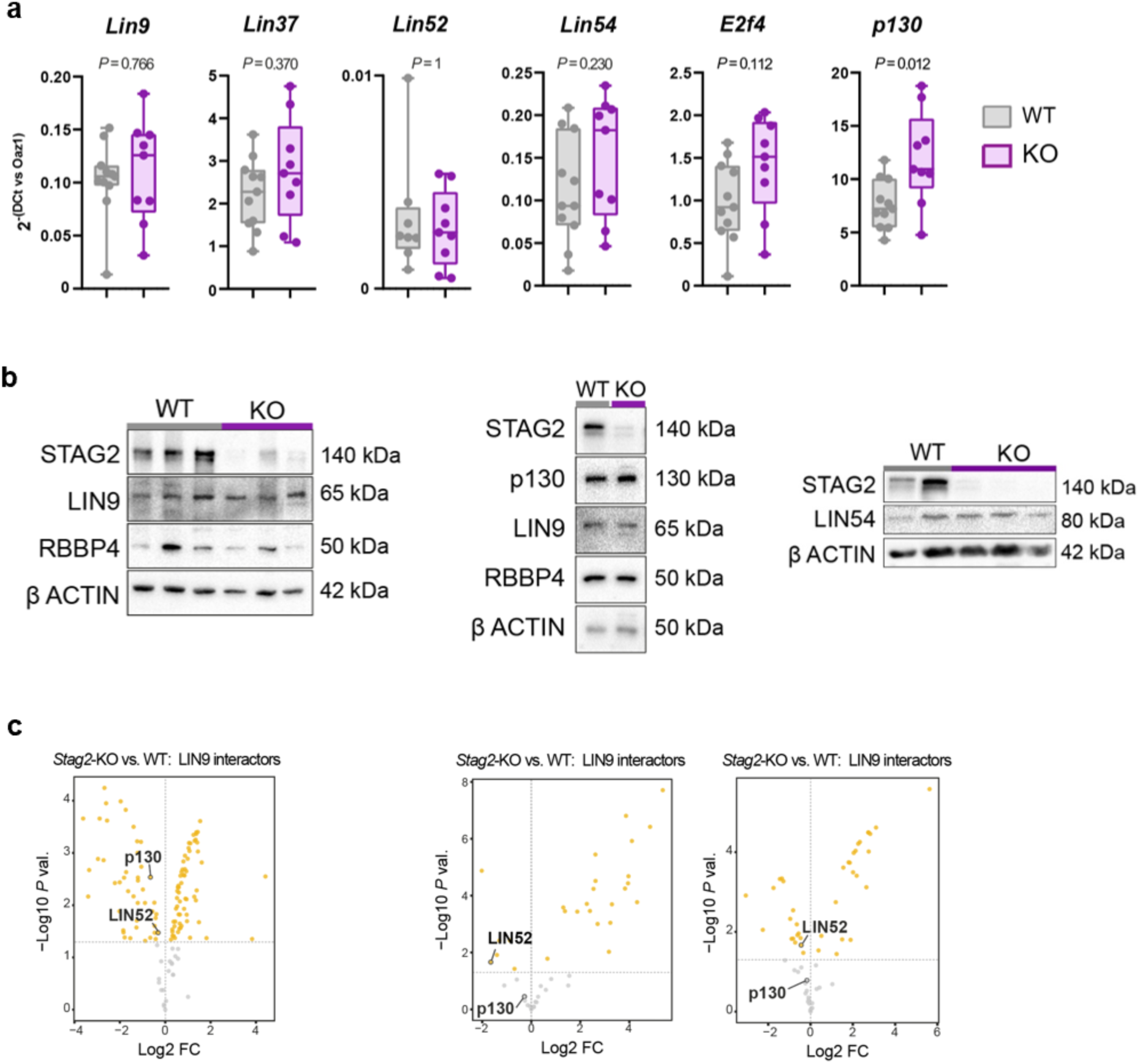
STAG2 loss in urothelial cells does not cause major changes in expression of DREAM components but reduces LIN52 and p130 binding to LIN9. **a,** RT-qPCR expression analysis of genes coding from DREAM components in WT and *Stag2*-KO mouse urothelium from *Trp63 ^CreERT^*^2^*;Stag2^lox/lox^*mice one week after inducing recombination. Data were normalized to *Oaz1* expression. Each dot represents one biological replicate. Statistical test: two-tailed Mann-Whitney. **b**, Western blot showing that STAG2 loss does not result in major differences in expression of DREAM components in *Stag2*-KO and WT quiescent organoids. Three representative experiments; the last one corresponding to the same experiment as in Suppl. Figure 5c. **c**, Volcano plots showing differential LIN9 interactors in *Stag2*-KO vs. WT quiescent organoids identified by IP-MS analysis in 3 independent experiments. LIN52 and p130 are highlighted. Coloured in yellow if *P* val < 0.05.

### *Stag2* inactivation in quiescent urothelial cells reduces genomic contacts associated with DREAM-binding sequences

To assess changes in chromatin looping, we performed Hi-C in *Stag2*-KO and WT quiescent organoids. We identified DNA loops at 25 kb resolution and found that *Stag2* inactivation resulted in a reduction of loop number (4,665 vs. 2,903 for WT and KO, respectively; example shown in **Fig. 7a**) and strength (**Fig. 7b**), and increased loop length (t-test *P* = 1E-175; **Fig. 7c**). Next, we defined sets of “common”, ‘lost’, and ‘gained’ interactions upon *Stag2* inactivation (**Fig. 7d**). Interestingly, common loops (1228 regions having the same bin for both anchors of the loop in both conditions) are also weaker in KO cells (Mann-Whitney *P* = 1E-41; **Fig. 7e**). Motif analysis of the regions engaged in these interactions revealed that CTCF was the top HOMER motif enriched in common interactions (*P* = 1E-4, adj. *P* = 0.005), but not among those lost nor gained (**Fig. 7f**). Importantly, MYB motifs (*de novo* HOMER analysis) were significantly over-represented exclusively in lost loop anchors (*P* = 1E-11) (**Fig. 7f**) while CHR motifs - the DREAM-binding sequences - were significantly enriched in the corresponding interacting regions (*P* = 1E-10) (**Fig. 7g**). We then intersected the collection of common, lost, and gained interactions with the set of STAG2-LIN54 shared binding sites identified in the Cut&Run experiments (**Fig. 5d,e**) and found that 52% of anchors occupied by STAG2 and DREAM were lost in *Stag2*-KO cells (**Fig. 7h**). This overlap and the enrichment of MYB motifs at lost loop anchors and of CHR motifs in their interacting regions suggest that *Stag2* inactivation alters DREAM binding sites and contacts. An exemplary affected gene was *Mybl2*, which codes for the MMB activator protein MYB-B. *Mybl2* is bound by STAG2 and DREAM at its promoter, belongs to a loop that is lost upon STAG2 loss (**Fig. 7i**), and is up-regulated in urothelial cells upon either *Stag2* or DREAM inactivation (**Fig. 2i,k,o**). These findings support the notion that STAG2 cooperates with DREAM by binding shared genomic sites to regulate gene expression via chromatin looping.

**Figure 7.**
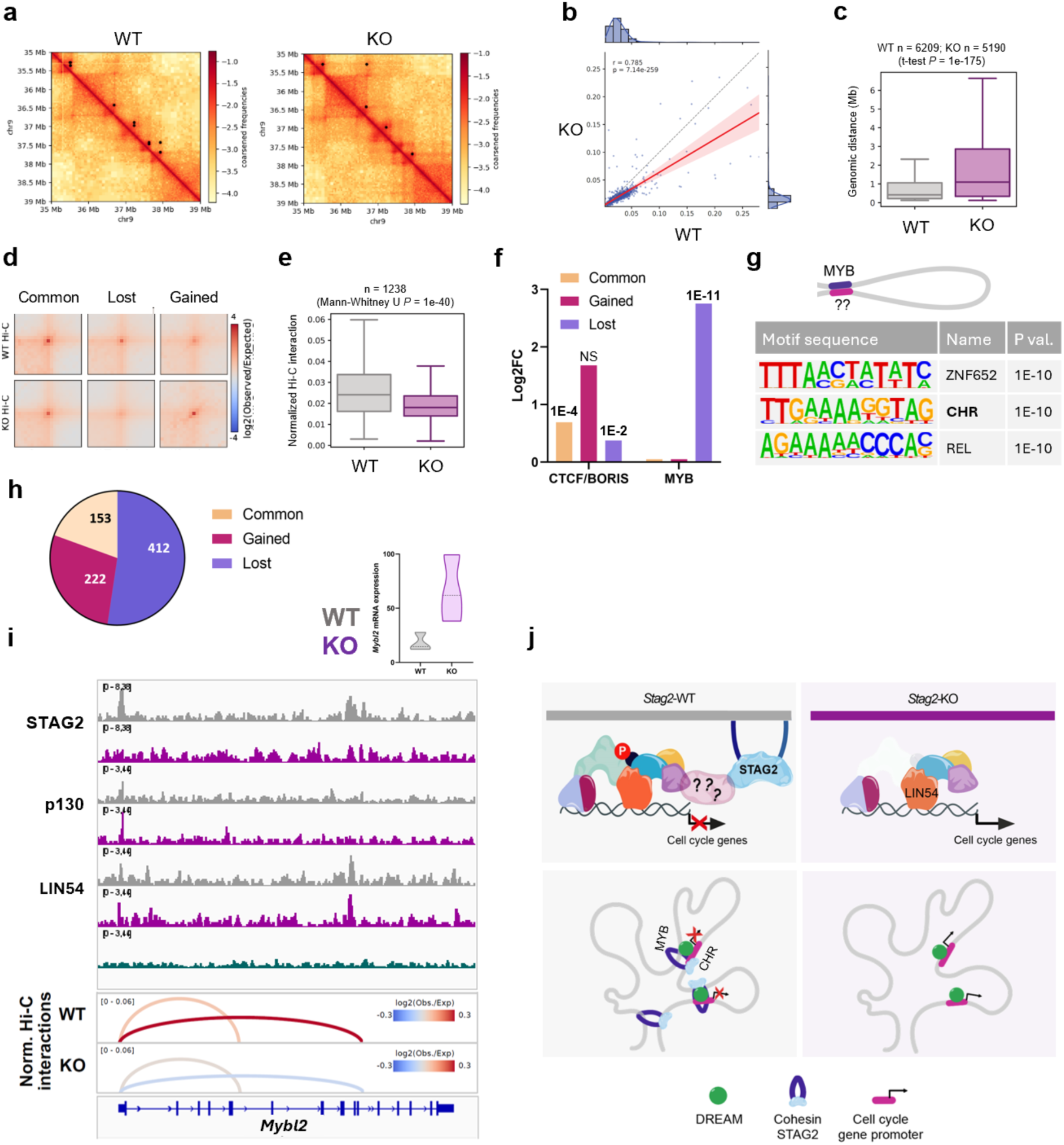
*Stag2* inactivation in quiescent urothelial cells leads to a reduced number and strength of chromatin interactions involving regions containing the MYB and CHR motifs. **a**, Heatmap showing exemplary reduced interactions upon *Stag2* inactivation. **b**, Reduced interaction strength upon *Stag2* inactivation. **c**, Increased loop length upon *Stag2* inactivation. **d**, Significantly gained and lost interactions. **e**, “Common” anchors are weaker upon *Stag2* inactivation. **f**, Motif analysis reveals that “Common” anchors are enriched in CTCF motifs (HOMER, known motifs) whereas “Lost” anchors are enriched in the MYB motif (HOMER, *de novo*), among others (only shown if *P* < 0.05), unlike “Gained”. The MYB motif is among the top 10 ones enriched in “Lost” anchors. Log2 of % of target sequences / % of background sequences with motif is shown; *P* val. for enrichment indicated on top of each bar. **g,** Scheme depicting a “Lost” region with MYB motifs and the corresponding interacting anchor, in which motifs are assessed. Top 3 motifs enriched. **h**, 52% of anchors containing STAG2-LIN54 shared binding sites are lost in *Stag2*-KO cells. **i**, Example of the *Mybl2* locus showing the combined genomic information in *Stag2*-WT and KO cells in grey and purple, respectively. Upper panel shows *Mybl2* mRNA expression, middle panels show Cut&Run data for STAG2, p130, and LIN54; and lower panel shows normalized Hi-C interactions. **j**, Model: In quiescent urothelial cells in homeostatic conditions, STAG2 cooperates with DREAM to repress cell cycle genes through binding common promoters and distal regions. In the absence of STAG2, DREAM-mediated repression is disrupted due to a reduced binding of LIN52, and possibly p130, to the MuvB core and to rewiring of chromatin interactions involving DREAM-binding motifs.

## Discussion

Genome sequencing studies have unveiled alterations in novel classes of genes that are significantly mutated in cancers, many of them through mechanisms (e.g., regulation of chromatin) different from those of classical oncogenes and tumour suppressors. In a study across 21 tumour types, only 12 genes were found to be significantly altered in >4 tumours, *STAG2* being one of them ^41^. *STAG2* is located on the X chromosome and mutations are predominantly loss-of-function ^3,19^. Despite many efforts, little is known about how STAG2 acts as a tumour suppressor.

Our previous studies of *STAG2* knockdown in RT112 bladder cancer cells showed a partial loss of urothelial identity with subtle changes in chromatin organization ^42^. Guided by the evidence that *STAG2* mutations are present in non-neoplastic urothelium of organ donors and the modest effects observed in RT112 cells, we turned to investigate the impact of *Stag2* genetic inactivation in normal urothelial cells using a variety of models including the conditional KO strain we described recently ^24^. While the sole inactivation of *Stag2* did not result in overt histological phenotypes in the urothelium, it conferred a primed state characterized by transient up-regulation of cell cycle genes and BrdU uptake, increased self-renewal capacity, and compromised activity of urothelial lineage identity programs. These effects may reflect a critical dependence on complete STAG2 depletion or on the cellular context (i.e., normal vs. transformed cells) and place *STAG2* as a novel class of tumour suppressors acting in the earliest steps of carcinogenesis, possibly through the expansion of mutant clones. While mosaicism has been well established in the haematopoietic system, little is known about clonal expansion in epithelial tissues ^14–16^.

Some of the effects of *STAG2* inactivation in the urothelium, such as increased proliferation, were transient whereas others were more sustained. Restoration of quiescence may rely on compensatory mechanisms implying - among others - the activity of pocket proteins or the DREAM complex ^6,43,44^. Additional growth promoting signals, including damage, inflammation, or oncogenic mutations, may overcome such compensatory mechanisms. Sustained effects include the partial loss of urothelial identity - with increased activity of “basal” programmes, increased *in vivo* regeneration potential and *in vitro* self-renewal capacity, and DNA damage. These features support that STAG2 loss confers memory to urothelial cells. We show that mutant, constitutively active, FGFR3 cooperates with STAG2 loss both *in vitro* and *in vivo* to promote cell proliferation and tumourigenesis, emphasizing the relevance of our findings to early urothelial carcinogenesis.

Using multiple omics strategies, we show that the transcriptional program activated upon STAG2 loss includes cell cycle genes located in chromatin-accessible regions in the quiescent urothelium. Our findings reveal that STAG2 cooperates with the DREAM complex to repress this programme by binding to shared genomic regions (**Fig. 7j**). Conversely, STAG2 loss compromises DREAM function by reducing LIN9 binding to LIN52 and - possibly - to p130. Both LIN52 and p130 are key to the assembly of the repressive DREAM complex at the promoter of cell cycle genes ^20,21^. Although we did not find evidence of a direct interaction of STAG2 and LIN9, STAG2 stabilizes the repressive DREAM complex and mediates short- and mid-range genome interactions, with 52% of anchors co-occupied by STAG2 and DREAM being lost in STAG2-null cells. The dual role of the DREAM complex in both repression and activation of cell cycle genes might explain why DREAM components have not been identified as cancer drivers ^45^. Yet, we describe here a novel mechanism through which the DREAM complex contributes to the homeostasis of tissues that are quiescent in the adult and to carcinogenesis.

We propose a model whereby STAG2 and DREAM bind to shared promoters and distal regulatory elements to control gene expression (**Fig. 7j**). The rewiring of chromatin interactions involving DREAM-binding motifs upon STAG2 loss provides a novel cancer-initiation mechanism that disrupts deep quiescence. Despite the major cell cycle regulatory capacity of DREAM, little is known about its role in adult tissue homeostasis *in vivo*. Our findings highlight that non-mutational mechanisms may contribute to DREAM’s dysfunction in slow cycling tissues.

## Acknowledgements

We thank María A. Blasco, Lars Dyrskjot, Luis C. Fernández, Jesús Gómez, Ana Losada, Juan Méndez, Sara Rodríguez, Eduardo Caleiras, Stéphanie Simon, and Gottfried Sjödahl for valuable contributions; the CNIO Molecular Imaging, Genome Editing, Genomics, Flow Cytometry, and Confocal Microscopy Units; members of the Epithelial Carcinogenesis Group for many valuable discussions; and Marcos Malumbres, Miguel Manzanares, Manuel Serrano, and Luciano Di Croce for comments to the manuscript.

## Author contributions

Conception and design: MR, EL, MAMR, and FXR

Financial support: EL and FXR

Administrative support: FXR

Provision of study materials: JX, FR, MJB, IBP and MLA

Collection and assembly of data: MR, EL, MNF, MM, EZ, MI, MN, JLMT, MMM, MAMR, and FXR

Data analysis and interpretation: MR, EL, MK, JMV, MNF, EZ, MI, OG, MAMR, and FXR Manuscript writing: MR, EL, MAMR, and FXR

Final approval of manuscript: MR, EL, MK, JMV, MNF, MM, EZ, MI, OG, JX, FR, MN, JLMT, MJB, IBP, MMM, MLA, MAMR, and FXR

Accountable for all aspects of the work: MR, EL, and FXR

## Methods

### Mice

The following mouse strains were used: *Stag2^lox^* ^24^, Tg.*UBC^CreERT2^* ^46^, *Trp63^CreERT2^* ^47^, *Tg.Upk3a^CreERT2^* [^48^, JAX #015855], *UpkII_hFGFR3-S249C* ^49^*, Dyrk1a^lox^* ^50^, *Rosa26_ACTB-tdTomato_EGFP* (*Rosa26^mT/mG^*) ^51^, and *Rosa26^Cas9^* ^32^. All crosses were established in a predominant *C57BL/6* background. Mice were maintained under specific pathogen free conditions. Experiments were performed using 8–20 week-old mice of both sexes, unless otherwise indicated. To activate CreERT2, TMX-containing food (Teklad CRD TAM^400^/CreER) was administered *ad libitum,* as indicated. Control mice carrying a wild-type (WT) *Stag2* (*Stag*2^+/+^) or a *CreERT2*^+/T^ ^or^ ^+/KI^ allele also received TMX.

BrdU was injected intraperitoneally (i.p.) (50 mg/kg) 1.5 h before sacrifice. Cyclophosphamide (CYC) (Merck Life Sciences S.L.U) was administered i.p. once (200 mg/kg) to induce urothelial damage. N-butyl-N-(4-hydroxybutyl)-nitrosamine (BBN) (0.025%) (Sigma) was administered in drinking water for 4 weeks, with the BBN solution replaced every 2-3 days. One week prior to starting BBN administration, mice received TMX (2mg/kg) i.p.

*hFGFR3-S249C;Stag2-KO* and *hFGFR3-S249C* mice underwent regular ultrasound scans (Vevo 3100, VisualSonics, Canada) with a RMV707b scan probe (40 MHz) under anesthesia. The entire abdominal area was scanned to detect structural alterations, including changes in echogenicity before and after bladder emptying.

Mice were sacrificed by CO_2_ inhalation. All experiments were carried out following guidelines for Ethical Conduct in the Care and Use of Animals as stated in The International Guiding Principles for Biomedical Research involving Animals, developed by the Council for International Organizations of Medical Sciences. Procedures were approved by the Institutional Animal Care and Use Committee, the Comité de Etica y Bienestar Animal, Instituto de Salud Carlos III (CEyBA, CBA 23_2020), and the General Guidance of the Environment of Madrid Community (PROEX 034.5/21).

### Establishment of NU1 cultures

Normal mouse bladder was digested with collagenase P (1.5 mg/ml in HBSS); after transfer to a 6-well plate, additional urothelial cells were detached using a scraper. Cells were collected by centrifugation, plated on collagen-coated plates, and cultured in Cascade Biologics Epilife Medium (Gibco M-Epi-500-CA) supplemented with 15% chelated fetal bovine serum (FBS), 5 ng/ml EGF, 1 X insulin/transferrin, 2.24×10^−11^ M T3 (3,3’,5-Triiodo-L-thyronine), 225 mg hydrocortisone, 5.3 ml cholera toxin, 0.3 mM CaCl₂, and 1 X penicillin/streptomycin (Gibco). NU1 cells were spontaneously immortalized through sequential passaging. To induce quiescence, cells were cultured in complete medium (CM) until confluence and then in CM supplemented with 1μM rosiglitazone (Merck) and 0.5 μM erlotinib.

### Establishment of mouse urothelial organoids

Mouse bladder urothelium was peeled and digested with collagenase P (Roche, 11213865001) (0.5 mg/mL in Hank’s Balanced Salt Solution (Gibco) at 37 °C with gentle shaking for 15 min. Organoids were established and maintained as described ^25^ (Table 1). Briefly, the urothelium was dissociated, enzymes were inactivated, the cell suspension was centrifuged and washed, and cells were embedded in Matrigel (Corning), on ice. Organoids were cultured in CM. For organoid expansion, cell recovery solution (Corning) was used; organoids were digested with 10 mg/mL dispase II. Single-cell suspensions were obtained by mechanical disruption with a 21G syringe. The pellet was washed and cells were resuspended in ice-cold Matrigel. To induce quiescence, organoids were cultured for 7-9 days in CM and then in growth factor-depleted medium (lacking WNT3A and RSPO1 conditioned medium, EGF, LY2157299 and noggin) for at least three days. To induce recombination, 2 mM 4-hydroxytamoxifen (4-OH-TMX) was added (Sigma) for three days. All experiments were performed using low-passage cultures (<10).

### Organoid-forming capacity and growth factor-dependency assays

Urothelial cell suspensions were obtained as described. Viable cells were isolated by sorting (FACS Aria IIU) and embedded in Matrigel (5,000 cells per 20-mL drop). To assess organoid-forming capacity, cultures were grown for 7 days in CM. Images were acquired in a wide-field platform (Leica Microsystems) and quantified using ImageJ.

Organoid quantification was performed using the Cellpose algorithm ^52^ by overlaying GFP and Tomato markers. Fluorescence intensity distribution of GFP and Tomato was calculated; organoids were classified as GFP+/- and Tomato+/- based on fluorescence in a predefined eroded area. Organoids with GFP^+^ or Tomato^+^ intensities above the mean plus two standard deviations were classified as “Full,” indicating high expression, while those below this threshold were labeled “Empty.” Fluorescence metrics, including maximum, minimum, mean, and integrated density, were computed for each marker.

### Immunofluorescence (IF) and immunohistochemical (IHC) analyses

Fresh tissues were fixed (4% buffered formaldehyde) and embedded in paraffin. Sectioning and H-E staining were done as described ^25^. For IHC, antigens were retrieved by boiling sections in citrate buffer. Endogenous peroxidase was inactivated with 3% H_2_O_2_ in methanol. Sections were incubated with blocking buffer (3% BSA in PBS/0.1% Triton X-100) for 1 h at room temperature (RT), then with primary antibodies (**Extended Data Table 2**) overnight at 4 °C. EnVision secondary antibodies (Dako) were added for 45 min at RT, reactions were developed with 3, 3’-diaminobenzidine, and nuclei were counterstained. Images were acquired in a Zeiss Imager A1 microscope and processed using the ZEISS-ZEN Software v3.8. For IF, sections were incubated in blocking buffer, primary antibodies (Table 2) were added overnight at 4 °C, and sections were next incubated with fluorochrome-conjugated secondary antibodies. Nuclei were counterstained with DAPI (1 μg/mL) and sections mounted with Prolong Gold Antifade Reagent (Life Technologies). Images were acquired using a Leica TCS SP5 Confocal microscope with an HC PL APO 20x/0.70 objective. Signal was quantified using QuPath Software v0.3.2 ^53^.

### Plasmids and lentiviral infections

Mission shRNAs (Sigma) were used for RNA interference. Lentiviruses were produced in HEK293T cells by FuGene-mediated transfection of the lentiviral construct together with packaging plasmids (psPAX2 and pCMV-VSVG). Medium was collected 48 h and 72 h after transfection, filtered, and stored frozen or added to NU1 cells in suspension in 5 mg/mL Polybrene (Sigma-Aldrich). The cells and viral particle mixture was spin-infected in and incubated for 1 h at 37 °C; cells were centrifuged and cultured.

### Cloning strategy to target *Lin54* by CRISPR/Cas9 in urothelial organoids

Single guide (sg) RNAs targeting the *Lin54* coding region were cloned into lentiGuide-Puro (AddGene 52963) ^54^. Products were used to transform One Shot Stbl3 cells. Plasmid DNA was isolated and purified. Lentiviral particles were produced as described above. Cells were cultured for 72 h and the supernatant was collected and filtered. Organoids were disaggregated into single cells and lentiviral particles were added with 5 mg/mL Polybrene); the mixture of cells and viral particles was spin-infected, incubated for 1 h at 37 °C, and cells were cultured as organoids as described. After 48 h, puromycin (1 μg/mL) was added.

### Protein extraction and western blotting

Organoids were recovered after removing Matrigel, pelleted, and proteins were solubilized with lysis buffer [2% SDS, 50 mM Tris-HCl (pH 8), 10 mM EDTA], sonicated, boiled, separated using 10% polyacrylamide SDS gel electrophoresis, and transferred onto nitrocellulose membranes. After blocking with 5% skim milk, they were incubated overnight with primary antibodies (**Extended Data Table 2**), washed, and incubated with HRP-conjugated secondary antibodies (Dako) for 45 min, and washed. Signal was detected using Immobilon Western HRP Chemiluminescent Substrate (Merk).

### Protein interaction assays

For co-immunoprecipitation, cells/organoids were pelleted and lysed in 0.05% IGEPAL® lysis buffer (50 mM Tris-HCl pH 7.4, 150 mM NaCl, 0.05% IGEPAL® CA-630) freshly supplemented with protease and phosphatase inhibitors. Proteins (0.5-1 mg) were incubated with antibodies (5 mg antibody/mg protein) overnight at 4 °C. Dynabeads™ Protein G (Invitrogen) (25 mL beads/sample) were added for 1 h at 4 °C and immune complexes were either eluted using 2X Laemmli buffer with 10% β-mercaptoethanol and boiled for 10 min for western blotting (IP-WB), or digested and processed for mass spectrometry (IP-MS).

After reduction and alkylation (by adding 6M urea, 50 mM TEAB pH 8.5, 15 mM TCEP, 50 mM CAA for 1 h in the dark at RT), proteins were digested with Lys-C (Wako) for 4h at RT (estimated enzyme:protein ratio, 1:50) followed by 200 ng trypsin (Promega) (estimated enzyme:protein ratio, 1:50), and incubated for 14 h at 37 °C. Resulting peptides were desalted using C18 stage-tips, speed-vac dried, and re-dissolved in 0.5% formic acid.

LC-MS/MS was performed by coupling an UltiMate 3000 RSLCnano LC system to either a Q Exactive Plus or Orbital Exploris 480 MS (Thermo Fisher Scientific). Peptides were loaded into a trap column (Acclaim™ PepMap™ 100 C18 LC Columns 5 µm, 20 mm length) for 3 min at a flow rate of 10 µl/min in 0.1% FA. Then, peptides were transferred to an EASY-Spray PepMap RSLC C18 column (Thermo) operated at 45 °C and separated using a 60 min effective gradient (buffer A: 0.1% FA; buffer B: 100% ACN, 0.1% FA) at a constant flow rate of 250 nL/min. The gradient ranged from 2% to 6% of buffer B in 2 min, from 6% to 33% B in 58 min, from 33% to 45% in 2 min, ending with 10 min at 98% B. Peptides were sprayed at 1.5 kV into the MS via the EASY-Spray source and the capillary temperature was set to 300°C. The mass spectrometer was operated in a data-independent acquisition (DIA) mode using 70,000 precursor resolution and 17,500 fragment resolution for Q Exactive Plus or 60,000 and 15,000 for Orbitrap Exploris 480. Ion peptides were fragmented using higher-energy collisional dissociation (HCD) with a normalized collision energy of 27 or 29.

The AGC target values were 3e6 for Full MS (maximum IT of 25 ms) and 1e6 for DIA MS/MS (maximum IT of 45 msec) for Q Exactive Plus or 300% for Full MS (maximum IT of 25 ms) and 1000% for DIA MS/MS (maximum IT of 22 msec) for Orbitrap Exploris 480. 8 m/z precursor isolation windows were used in a staggered-window pattern from 396.43 to 1004.70 m/z. A precursor spectrum was interspersed every 75 DIA spectra. The scan range of the precursor spectra was 390-1000 mz.

Raw files were processed with DIA-NN 1.8.2 using the library-free setting against a mouse protein database (UniProtKB/Swiss-Prot, one protein sequence per gene, 21,990 sequences). Precursor m/z range was set from 390 to 1010, and all other settings were set at their default values. A protein group intensity table was obtained by summing the precursor quantity values from the table report.tsv after filtering for a Lib.Q.Value and Lib.PG.Q.Value < 0.01. Prostar 1.26.4 ^55^ was used for further statistical analysis. Briefly, protein intensity was normalized using the median intensity. Proteins with 50% or less valid values in at least one experimental condition were filtered out. Missing values were imputed using the algorithms SLSA ^56^ for partially observed values and DetQuantile for values missing on an entire condition. Differential analysis was done using the empirical Bayes statistics Limma. Proteins with a *P* val. < 0.05 and a log2 ratio >2 were defined as potential interactors. For the *Stag2*-KO vs. WT comparison, potential interactor intensities were normalized using bait intensity. For every comparison, the FDR was estimated to be below 6% by Benjamini-Hochberg.

### RNA extraction and real-time quantitative PCR

Total RNA was extracted using a ReliaPrep^TM^ System Kit (Promega), according to manufacturer’s instructions. Reverse transcription and PCR amplification were performed as described ^25^. Gene-specific expression was normalized to *Oaz1* using the ΔΔCt method. Primers used are indicated in **Extended Data Table 3**.

### RNA-Seq and data processing

RNA was extracted (3 replicates per experimental condition) and quality was assessed using an Agilent 2100 Bioanalyzer. cDNA sequencing libraries were prepared following the manufacturer’s recommendations (Lexogen) and sequenced in an Ilumina NextSeq 550 instrument. RNA-Seq data were processed as described ^57^ using default settings. Briefly, FastQC was used to assess read quality and FastqScreen to evaluate cross-contamination. Raw reads were preprocessed using BBDuk Software to trim contaminant sequences. Reads were aligned to the mouse reference genome (mm10) using Bowtie. Gene counts were generated from alignment maps using HTSeq-count. Count normalization and differential expression analysis were performed using DESeq2 ^58^ in R Studio. Statistical significance was set at adjusted *P* value (adj. *P* val.) < 0.05. Batch or experimental biases were corrected, if needed, by adjusting the design formula in DESeq2.

Pathway enrichment analysis was conducted with the online software Enrichr (https://maayanlab.cloud/Enrichr/) using the BioPlanet 2019 database and Gene Set Enrichment Analysis (GSEA) was performed using the GSEA Software v4.3.2. Promoter motif analysis was run with Hypergeometric Optimization of Motif EnRichment (HOMER) software findMotifs.pl ^59^ using default settings, searching for motifs from −400 bp to +100 bp relative to the transcriptional start site (TSS).

### RNA-Seq analysis of published transcriptomic data

The following public RNA-Seq data were used: 21 normal human bladders from the Genotype-Tissue Expression (GTEx) database (https://www.gtexportal.org/home/index.html) ^60^ and 320 low-grade NMIBC from the UROMOL project ^61,62^. Data was analyzed with the R Software v4.3.0 or higher using RStudio. DESeq2 was used to perform differential expression analyses. Significant DEGs were those with adj. *P* val. < 0.05. GSEA was performed using the GSEA v4.3.2 software.

### ATAC-Seq

ATAC-Seq was conducted as described, in triplicates ^63^. Briefly, 50,000 viable GFP^+^ cells were isolated by FACS. Cells were pelleted and resuspended in 50 mL of cold Omni-ATAC lysis buffer. Nuclei were washed with Omni-ATAC wash buffer pelleted and resuspended in transposition mix. The reaction mix was incubated for 30 min at 37 °C. DNA was eluted using the MinElute PCR Purification Kit (Qiagen) following manufacturer’s instructions. ATAC-Seq libraries were prepared using the NEBNext 2 X PCR Master Mix (NEB) as described ^64^ and purified using 1.8 X AMPure XP beads (Beckman Coulter). Libraries were sequenced in an Illumina NextSeq 550 instrument. Paired-end sequencing data was analyzed using the ENCODE ATAC-Seq pipeline (https://github.com/ENCODE-DCC/atac-seq-pipeline) with default parameters. Briefly, ^65^reads were aligned to the reference genome (mm10) using Bowtie2, and unmapped reads, reads aligned to mitochondrial DNA, and PCR duplicates were discarded. MACS2 was used to define accessible regions. Peaks with *P* < 0.01 were used for analysis. Blacklisted regions were filtered out and differentially accessible regions (DARs) between experimental conditions were identified with DiffBind in R studio. Accessible regions found in at least 2 out of 3 replicates were considered. Significant DARs were those with FDR < 0.05. Peaks of interest were annotated with the HOMER annotatePeaks.pl function using the mm10 genome. Promoter regions were defined as the genomic space spanning from −1 kb to +1 kb relative to the TSS. Motif analysis was performed with HOMER findMotifsGenome.pl ^59^using the argument “-size given”.

### ChIP-Seq and data processing in mouse cells

ChIP-Seq was performed as previously described, in duplicates ^42^. Briefly, cells (4 × 10^7^) were washed with PBS, trypsinized, and chromatin was crosslinked for 15 min at RT. After quenching with 0.125 M glycine, cells were washed and lysed. Chromatin was sonicated in a Covaris instrument for 30 min (20% duty cycle; 6% intensity; 200 cycle), yielding DNA fragments of 200–300 bp. Chromatin was diluted and samples pre-cleared with a mix of BSA-blocked protein A/G agarose beads for 2 h at 4°C and centrifuged. The supernatant was divided into aliquots (550 μg/assay) and incubated with 25 μg of antibody (**Extended Data Table 2**; STAG1, STAG2, CTCF, non-related IgG) overnight at 4 °C on a rotating platform. Next, protein A/G agarose beads (Santa Cruz) were added for 2 h at 4 °C on a rotating platform and beads were sequentially washed with low-salt wash buffer, high-salt wash buffer, LiCl wash buffer, and TE 1X, as described ^42^. DNA was eluted (1% SDS, 0.1 M NaHCO_3_) and cross-linking was reversed by overnight incubation with 5M NaCl in dilution buffer at 65 °C. RNA and proteins were digested with 10 mg/ml RNase (Life Technologies) and 10 mg/ml of proteinase K (ITW reagents). DNA was purified by phenol–chloroform extraction.

For library preparation, 5 ng of DNA per condition were used. Samples underwent sequential enzymatic treatments of end-repair, dA-tailing, and ligation to adapters with ‘NEBNext Ultra II DNA Library Prep Kit for Illumina’ (New England BioLabs, ref. E7645). Adapter-ligated libraries were completed by limited-cycle PCR and extracted with a single double-sided SPRI size selection. The resulting average fragment size was 370 bp, from which 120 bp corresponded to adaptor sequences. Libraries were applied to an Illumina flow cell for cluster generation and sequenced on an Illumina NextSeq 500.

ChIP-Seq data were analyzed using the Rubioseq pipeline ^66^. Briefly, reads were aligned to the reference genome (mm10) using BWA ^67^ and duplicates removed with Picard. Genomic regions were called using MACS2. Consensus peak sets per experimental condition were generated using the mergePeaks function of HOMER. For STAG1 and STAG2 ChIP-Seq, significant peaks were defined based on the following criteria: FDR < 0.05, fold enrichment vs. input > 10, and pileup > 30. Motif enrichment analysis was performed using the findMotifsGenome.pl function of HOMER. Peak annotation and pathway enrichment analyses were performed as described.

### Cut&Run

Urothelial cells (approx. 500,000 cells for each experimental condition) were analyzed (2-3 replicates per experimental condition). Cut&Run was performed as described^68^. Briefly, BioMag®Plus concanavalin A-coated magnetic beads (Polysciences) were activated using binding buffer. Cells were washed in BSA-coated tubes using wash buffer. Cells were immobilized on activated concanavalin A-coated magnetic beads for 10 min at RT. The suspension was incubated with the corresponding antibody (**Extended Data Table 2**) in wash buffer supplemented with 2 mM EDTA and 0.025% digitonin, for cell permeabilization, overnight at 4 °C. The next day, cells were washed and incubated with 700 ng/mL pAG-MNase in wash buffer containing 0.025% digitonin for 1 h at 4 °C. Digestion was performed by adding 100 mM Ca^2+^ for 1 h at 0 °C and stopped with stop buffer containing 2 pg/mL heterologous *S. cerevisiae* as spike-in genome. To release DNA-protein complexes, cells were incubated for 30 min at 37 °C. DNA fragments were extracted from the supernatant using 0.1% SDS and 0.1 mg/mL proteinase K and incubating for 10 min at 70 °C. DNA was purified by phenol/chloroform extraction followed by ethanol precipitation, air-dried, and resuspended in nuclease-frese water. Libraries were prepared as described ^69,70^ and applied to an Illumina flow cell for cluster generation and sequenced on an Illumina NextSeq 550 instrument by following manufacturer’s recommendations.

Cut&Run sequencing data was analysed using the Cut&Run Nextflow pipeline ^70^. Briefly, reads were trimmed and aligned to mm10 reference genome and to *S. cerevisiae* as spike-in genome using Bowtie2; duplicates were removed using Picard. Peak calling was performed using MACS2 with pMNase-treated chromatin as control samples. FDR-significant peaks <0.05 were used for analysis. Peaks were merged using the mergePeaks HOMER function with the argument “-size given”. Promoter regions were defined as the genomic space spanning from −1 kb to +1 kb relative to the TSS. Peak annotation, pathway enrichment, motif enrichment, and differential binding analyses were performed as described.

### In situ Hi-C and data analysis

In situ Hi-C was performed as described, in duplicates (Rao et al., 2014). Organoid suspensions were plated as 10 mL in Matrigel drops (12,500 cells/drop), maintained in CM for 5 days, and then cultured for 3 days in GF-depleted medium with 4-OH-TMX. After crosslinking (1% (w/v) methanol-free formaldehyde 10 min RT 65 rpm), the reaction was quenched with 0.125 M glycine (5 min RT 65 rpm) followed by 30 min at 4 °C. Organoid suspensions were collected on ice, centrifuged (3 min, 300 g at 4 °C), washed with cold PBS, dissociated with 0.25% Trypsin-EDTA, and disaggregated for 3 min. Cells were centrifuged, washed, flash-frozen, and stored at −80 °C.

Cells were lysed in lysis buffer for 30 min on ice. Lysates were centrifuged and pellets were washed with 1x NEBuffer 2, resuspended in NEBuffer 2/0.5% SDS, and incubated for 10 min at 65 °C at 350 rpm. SDS was quenched by adding 400 mL 1x NEBuffer 2 and 120 mL 10% Triton X-100 (Merck, 648466), followed by incubation at 37 °C for 30 min at 300 rpm. Nuclei were washed with 1x NEBuffer 2, pelleted, and resuspended in 300 mL 1x NEBuffer 2 and 400 Units of MboI (NEB). Chromatin was digested overnight at 37 °C 300 rpm. Digestion efficacy was assessed using Tapestation (Agilent Technologies).

#### Biotinylation and ligation

Samples were centrifuged and pellets resuspended with reparation mix (30 mL 10x NEBuffer 2, 1.5 mL 10 mM dCTP (Roche), 1.5 mL 10 mM dTTP (Roche, 11969064001), 1.5 mL 10 mM dGTP (Roche), 37.5 mL 0.4 mM biotin-dATP (ThermoFisher), 4.5 mL 50,000 U/ml Klenow (NEB) and water up to 300 mL). The mix was incubated for 45 min at 37 °C, followed by 65 °C for 10 min. Samples were next centrifuged at 16 °C 300 rpm and pellets incubated overnight in ligation mix [120 mL 10x NEB T4 ligase buffer (NEB), 100 mL 10% Triton X-100, 6 mL 20 mg/ml recombinant albumin (NEB), 5 mL 2,000,000 U/ml T4 DNA Ligase (NEB) and water up to 1.2 ml].

#### Cross-link reversion and purification

Nuclei were resuspended with 400 mL 1x NEBuffer 2 and 10 mL 10 mg/ml RNase A (ThermoFisher) and incubated for 15 min at 37 °C. Proteinase K was added and incubated for 6 h at 65 °C at 350 rpm. DNA purification was performed with 1x volume AMPure beads.

#### Shearing and library preparation for sequencing

DNA (1 mg) was sheared in a Bioruptor Pico (Diagenode) to 280-330 bp fragments in cycles of 20 sec ON and 60 sec OFF (6-8 cycles). The final volume was adjusted to 200 mL with water and pull-down of the biotin tagged Hi-C fragments was done by mixing with 200 mL washed Dynabeads MyOne Streptavidin T1 beads (ThermoFisher) and incubating for 30 min with rotation at RT. Libraries were prepared with NEBNext End Repair (NEB) and dA-Tailing Module (NEB) following manufacturer’s recommendations and washing twice with 1x BB buffer (5 mM Tris-HCl pH7.5, 0.5 mM EDTA, 1 M NaCl) between reactions. Beads were resuspended in 50 mL Adapters Ligation mix (10 mL 5x Quick Ligation Reaction Buffer (NEB), 2.5 mL NEBNext Adapter (NEB), 2 mL 2,000,000 U/ml T4 DNA Ligase (NEB) and nuclease-free water up to 50 mL) and incubated for 15 min at RT. Next, 3 mL USER (NEB) was added and incubated for 15 min at 37 °C. Finally, beads were resuspended in 20 mL water and all volume was used for the final amplification with 5 mL UDIs (NEB) and 25 mL NEBNext High-Fidelity 2x PCR Master Mix (NEB). Amplification was done 30 sec at 98 °C, 8 cycles of 10 sec at 98 °C, 30 sec at 65 °C and 30 sec at 72 °C and a final extension of 5 min at 72 °C. The amplified DNA fragments were purified with 0.9x volume AMPure beads, the library profile was assessed in Tapestation and sequenced with Illumina paired end sequencing up to 500 M reads.

#### Hi-C mapping and analysis

Hi-C data were processed using an in-house pipeline based on TADbit ^71^. Reads were mapped as described ^72^. Read ends were mapped to the reference mouse genome (GRCm38/mm10). TADbit filtering module was used to remove noninformative contacts and create contact matrices ^71^. PCR duplicates were removed and the Hi-C filters applied corresponded to potential nondigested fragments (extra-dangling ends), non-ligated fragments (dangling ends), self-circles, and random breaks. Contact matrices next were normalized for sequencing depth and genomic biases using cooler balance^73^. Further analyses were performed by cooltools ^74^ and in-house scripts at variable genomic resolutions. Motif analysis was run with HOMER using default settings. Overlap between loops and Cut&Run peaks was assessed using HOMER mergePeaks.

### Statistical analyses

Statistical analyses were performed using GraphPad Prism Software v8.0.1 or R Software v4.3.0 or higher using RStudio. The statistical test applied in each experiment is indicated in the figure legend. D’Agostino test was used to assess data normality and statistical significance was calculated using a two-tailed t-test for normally distributed data and a Mann-Whitney test otherwise. Fisher’s exact test was used to assess statistical differences to examine the significance of the association of categorical data. *P* < 0.05 was considered significant. Data are presented as means and standard deviation, or as box plots with median, IQR, and whiskers extending to the minimum and maximum values. The ROUT method (Robust Regression and Outlier Removal) was applied to identify outliers in qPCR experiments and in the quantification of positive cells by IF/IHC when outliers were suspected, using a maximum false discovery rate of 1%.

### Data accessibility

Proteomics data can be found at ProteomeXchange under the Project accession number: PXD060155.

## Reviewer access details

**Proteomic data**: Log in to the PRIDE website using the following details:

Project accession: PXD060155

Token: 5NpZLh5CUw83

Alternatively, reviewers can access the dataset by logging in to the PRIDE website using the following account details:

Username

Password: HcAFAaiRnCNA

**Genomic data**: RNA-Seq data have been deposited at the Gene Expression Omnibus repository (NU1 cells, GSE288619; normal mouse urothelium, GSE288629); ChIP-Seq, Cut&Run, and HiC data are being deposited at the Gene Expression Omnibus repository and accession number is awaited.

## Extended Data Supplementary Datasets

**Suppl. Dataset 1.** RNA-Seq data from *Stag2*-KO vs. WT peeled urothelium (1 week and 1 month) and *Stag2*-KD vs. WT proliferating and quiescent NU1 cells.

**Suppl. Dataset 2.** Interactors identified in IP-MS for endogenous LIN9 in *Stag2*-KO and WT quiescent urothelial organoids.

## Supplementary Tables

**Supplementary Table 1.** List of reagents used for organoid culture.

| Reagent | Concentration | Source, Ref. |
| --- | --- | --- |
| Advanced DMEM/F12 medium | 1 X | Gibco, 12634028 |
| B27 (without Vitamin A) | 1 X | Gibco, 12587010 |
| Bovine serum albumin | 1% | Sigma-Aldrich, A7906 |
| Cell recovery solution | 1 X | Corning, 45354253 |
| Charcoal-stripped fetal bovine serum | 50% | Gibco, 12676029 |
| Collagenase P | 0.5 mg/mL | Roche, 11213865001 |
| Dispase II | 10 mg/mL | Gibco, 17105041 |
| EDTA | 2 mM | Sigma-Aldrich, E6758 |
| GlutaMAX | 1 X | Gibco, 35050038 |
| Hank's balanced salt solution | 1 X | Gibco, 14025050 |
| HEPES | 1 X | Gibco, 15630056 |
| LY2157299 | 1 mM | AxonChem, Axon 1491 |
| Matrigel | 1 X | Corning, 356231 |
| N2 | 1 X | Gibco, 17502048 |
| N-acetylcysteine | 1 mM | Sigma-Aldrich, A9165 |
| Penicillin/streptomycin | 1 X | Gibco, 15070-063 |
| Recombinant human noggin | 100 ng/mL | Peptotech, 120-10C |
| Recombinant mouse EGF | 50 ng/mL | Life Technologies, PMG8041 |
| RSPO1 conditioned medium | 5% | In-house production |
| WNT3A conditioned medium | 50% | In-house production |
| Y27632 | 10 mM | Selleck Chemicals, S1049 |
| Trypsin-EDTA | 1 X | Gibco, 25200056 |

**Supplementary Table 2.**
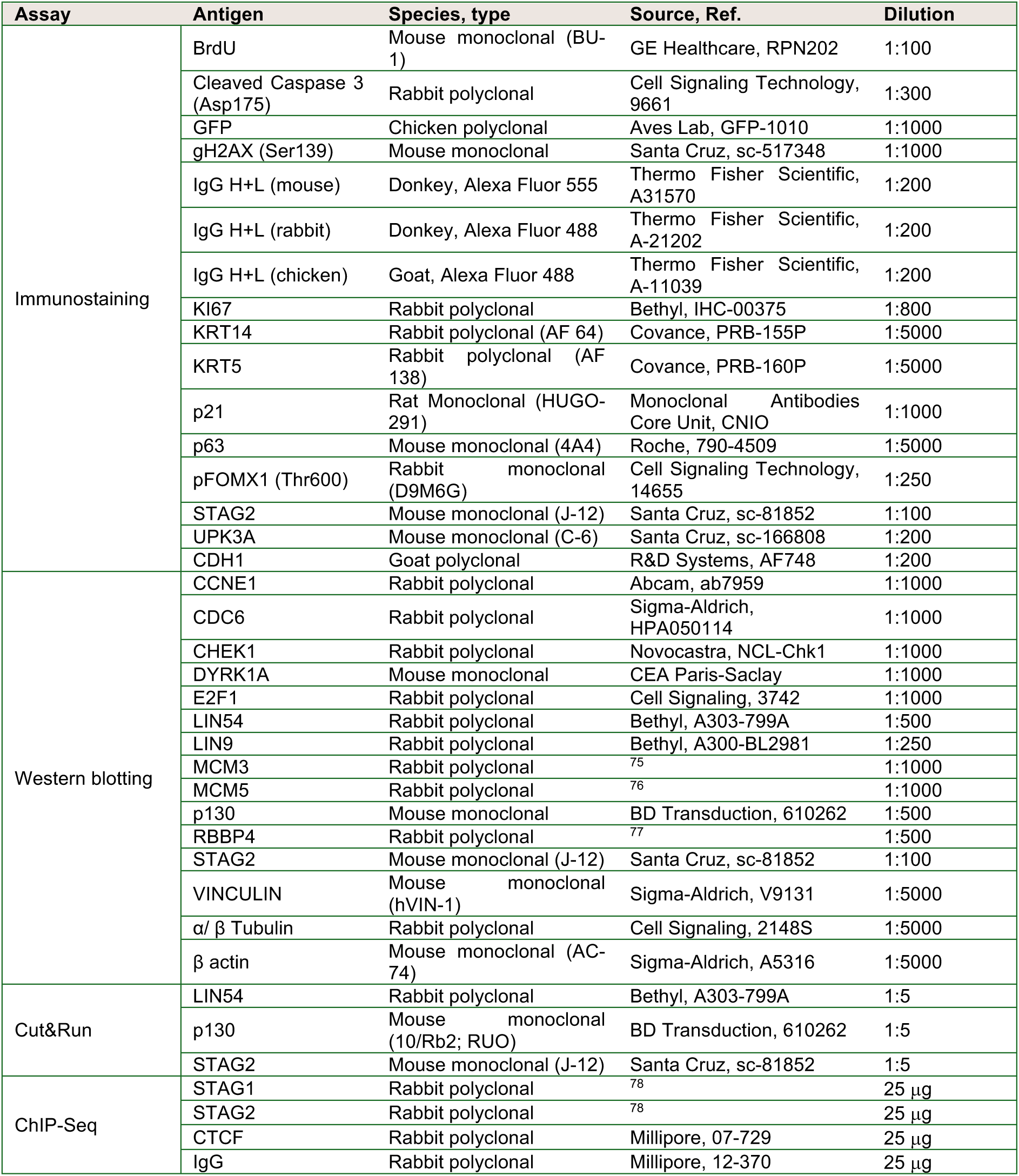
List of antibodies used for immunostaining. Antigen names are alphabetically ordered and grouped by experimental assay.

**Supplementary Table 3.**
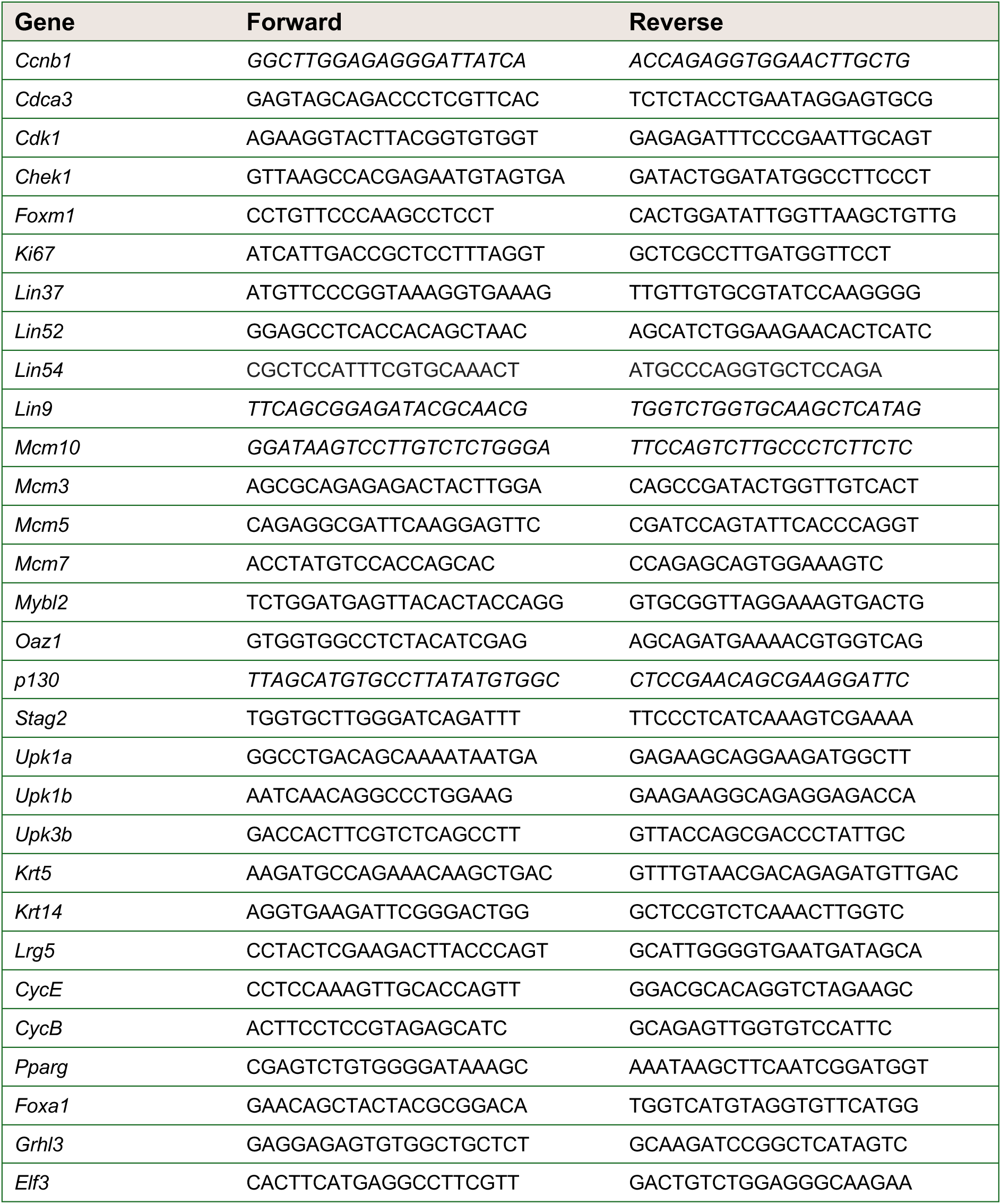
List of primers used for RT-qPCR.

